# Multifaceted N-degron recognition and ubiquitylation by GID/CTLH E3 ligases

**DOI:** 10.1101/2021.09.03.458554

**Authors:** Jakub Chrustowicz, Dawafuti Sherpa, Joan Teyra, Mun Siong Loke, Grzegorz Popowicz, Jerome Basquin, Michael Sattler, J. Rajan Prabu, Sachdev S. Sidhu, Brenda A. Schulman

## Abstract

N-degron E3 ubiquitin ligases recognize specific residues at the N-termini of substrates. Although molecular details of N-degron recognition are known for several E3 ligases, the range of N-terminal motifs that can bind a given E3 substrate binding domain remains unclear. Here, studying the Gid4 and Gid10 substrate receptor subunits of yeast “GID”/human “CTLH” multiprotein E3 ligases, whose known substrates bear N-terminal prolines, we discovered capacity for high-affinity binding to diverse N-terminal sequences determined in part by context. Screening of phage displaying peptide libraries with exposed N-termini identified novel consensus motifs with non-Pro N-terminal residues distinctly binding Gid4 or Gid10 with high affinity. Structural data reveal that flexible loops in Gid4 and Gid10 conform to complementary folds of diverse interacting peptide sequences. Together with analysis of endogenous substrate degrons, the data show that degron identity, substrate domains harboring targeted lysines, and varying E3 ligase higher-order assemblies combinatorially determine efficiency of ubiquitylation and degradation.

## INTRODUCTION

Specificity of ubiquitylation depends on E3 ligases recognizing motifs, termed “degrons”, in substrates to be modified. The first such motif to be identified was the N-terminal sequence - now called N-degron [1] - in substrates of the yeast E3 ligase Ubr1 [2–4]. Subsequently, several E3 ligases in different families were discovered to recognize protein N-termini as degrons. Higher eukaryotes have one HECT-type and several RING-family E3s with “Ubr” domains homologous to those in yeast Ubr1 that either have been shown to or are presumed to recognize distinct N-terminal sequences [5, 6]. Other N-degron-recognizing ubiquitin ligases were identified either through characterizing substrate sequences mediating E3-binding [7, 8], or through systematic genetic screens matching human protein N-terminal sequences with E3 ligases [9]. Some of the best-studied pathways recognize sequences with an N-terminal Arg [2], Pro [7, 10] or Gly [9, 11] (termed Arg/N-degron, Pro/N-degron or Gly/N-degron, respectively), or acetylated N-terminus (Ac/N-degron) [12–15].

An N-degron-recognizing E3 of emerging importance is a suite of related multiprotein complexes termed “GID” in budding yeast (named due to mutations causing *g*lucose-*i*nduced degradation *d*eficiency of fructose-1,6-bisphosphatase, Fbp1) [8, 16–20] or “CTLH” in higher eukaryotes (named due to preponderance of subunits containing CTLH motifs) [21]. The yeast GID E3 mediates degradation of gluconeogenic enzymes Fbp1, Mdh2 and Icl1 during recovery from carbon starvation [8]. The GID E3 recognizes the N-terminal Pro in these substrates generated by cleavage of the initiator methionine [7, 8]. In higher eukaryotes, corresponding CTLH complexes are involved in diverse biological processes including erythropoiesis, organ development, embryogenesis, and cell division [22–32]. However, the mechanistic roles of CTLH-mediated ubiquitylation in these pathways remain largely mysterious.

Recent genetic, biochemical and structural studies have revealed that the GID E3 is not a singular complex. Rather a core GID^Ant^ complex (comprising Gid1, Gid5, Gid8, Gid2, Gid9 subunits) essentially anticipates shifts in environmental conditions that stimulate expression of interchangeable and mutually exclusive substrate-binding receptors – Gid4 (termed “yGid4” for yeast Gid4 hereafter) [17, 33, 34], Gid10 (yGid10 hereafter) [34–36] and Gid11 (yGid11 hereafter) [37]. Whereas yGid4 is expressed after glucose has been restored to carbon-starved yeast, yGid10 and yGid11 are upregulated upon other environmental perturbations including heat shock, osmotic stress as well as carbon, nitrogen and amino acid starvation. The resultant E3 complexes, GID^SR4^, GID^SR10^, and GID^SR11^ (where SR# refers to Gid substrate receptor), recognize distinct N-terminal sequences of their substrates [7, 8, 34, 35, 37]. In addition, another subunit, Gid7, can drive supramolecular assembly of two GID^SR4^ units into a complex named Chelator-GID^SR4^ to reflect its resemblance to an organometallic chelate capturing a smaller ligand through multiple contacts [38]. The cryo EM structure of a Chelator-GID^SR4^ complex with Fbp1 showed two opposing Gid4 molecules avidly binding N-degrons from different Fbp1 protomers. As such, Fbp1 is encapsulated within the center of the oval-shaped Chelator-GID^SR4^. This assembly positions functionally-relevant target lysines from multiple Fbp1 protomers adjacent to two Chelator-GID^SR4^ catalytic centers.

The molecular details of GID/CTLH recognition of Pro/N-degrons were initially revealed from crystal structures of human Gid4 (referred to as hGid4 hereafter) bound to peptides with N-terminal prolines [10]. Although Pro/N-degron substrates of the CTLH E3 remain unknown, hGid4 is suitably well-behaved for biophysical and structural characterization, whereas yGid4 has limited solubility on its own [10]. Previously, the sequence PGLWKS was identified as binding hGid4 with highest affinity amongst all sequences tested, with a K_D_ in the low micromolar range [10]. The crystallized peptide-binding region of hGid4, which superimposes with the substrate-binding domains of yGid4 and yGid10 in GID^SR4^ and GID^SR10^, adopts an 8-stranded β-barrel with a central tunnel that binds the N-terminus of a peptide, or of the intrinsically-disordered N-terminal degron sequence of a substrate [10, 34, 36, 38, 39]. Loops between β-strands at the edge of the barrel bind residues downstream of the peptide’s N-terminus. Interestingly, although GID^SR4^ was originally thought to exclusively bind peptides with an N-terminal Pro, hGid4 can also bind peptides with non-Pro hydrophobic N-termini such as Ile or Leu, albeit with at best ≈8-fold lower affinity [39]. Furthermore, yGid11 is thought to use a distinct structure to recognize substrate Thr/N-degrons [37]. Collectively, these findings suggested that the landscape of GID/CTLH E3 substrates can extend beyond Pro/N-degron motifs.

Here, phage display screening identified peptides with non-Pro N-termini that not only bind hGid4, yGid4 and yGid10, but do so with comparable or higher affinity than the previously identified Pro-initiating sequences including Pro/N-degrons of ubiquitylation substrates. Structural data reveal that loops in GID/CTLH substrate-binding domains adopt conformations complementary to partner peptide sequences downstream of the N-terminus. Thus, sequence context is a determinant of N-terminal recognition by GID/CTLH substrate-binding domains. In the context of natural substrates recognized by yGid4, not only the degron but also the associated domain harboring targeted lysine contribute to ubiquitylation by the core GID^SR4^ and its superassembly.

## RESULTS

### hGid4 can bind peptides with a range of N-terminal sequences

We took advantage of the amenability of hGid4 to biophysical characterization to further characterize features of the PGLWKS sequence mediating interactions. To assess the importance of peptide length beyond the N-terminus, we examined chemical shift perturbations (CSPs) in 2D ^1^H, ^15^N-HSQC NMR spectra of [^15^N]-labeled hGid4 mixed with the amino acid Pro, a Pro-Gly dipeptide, or the PGLWKS peptide (Fig. 1A). Although prior studies emphasized the importance of an N-terminal Pro [10, 39], Pro alone only minimally influenced the spectrum. The Pro-Gly dipeptide elicited stronger CSPs, presumably due to the peptide bond directly interacting with hGid4, and suppressing repulsion by burying the negatively charged carboxylate of a single Pro in a hydrophobic environment (Fig. S1A). The PGLWKS peptide showed the greatest CSPs and binding kinetics in the slow exchange regime at the NMR chemical shift time scale, indicating tight binding, and, therefore, importance of downstream residues.

**Figure 1:**
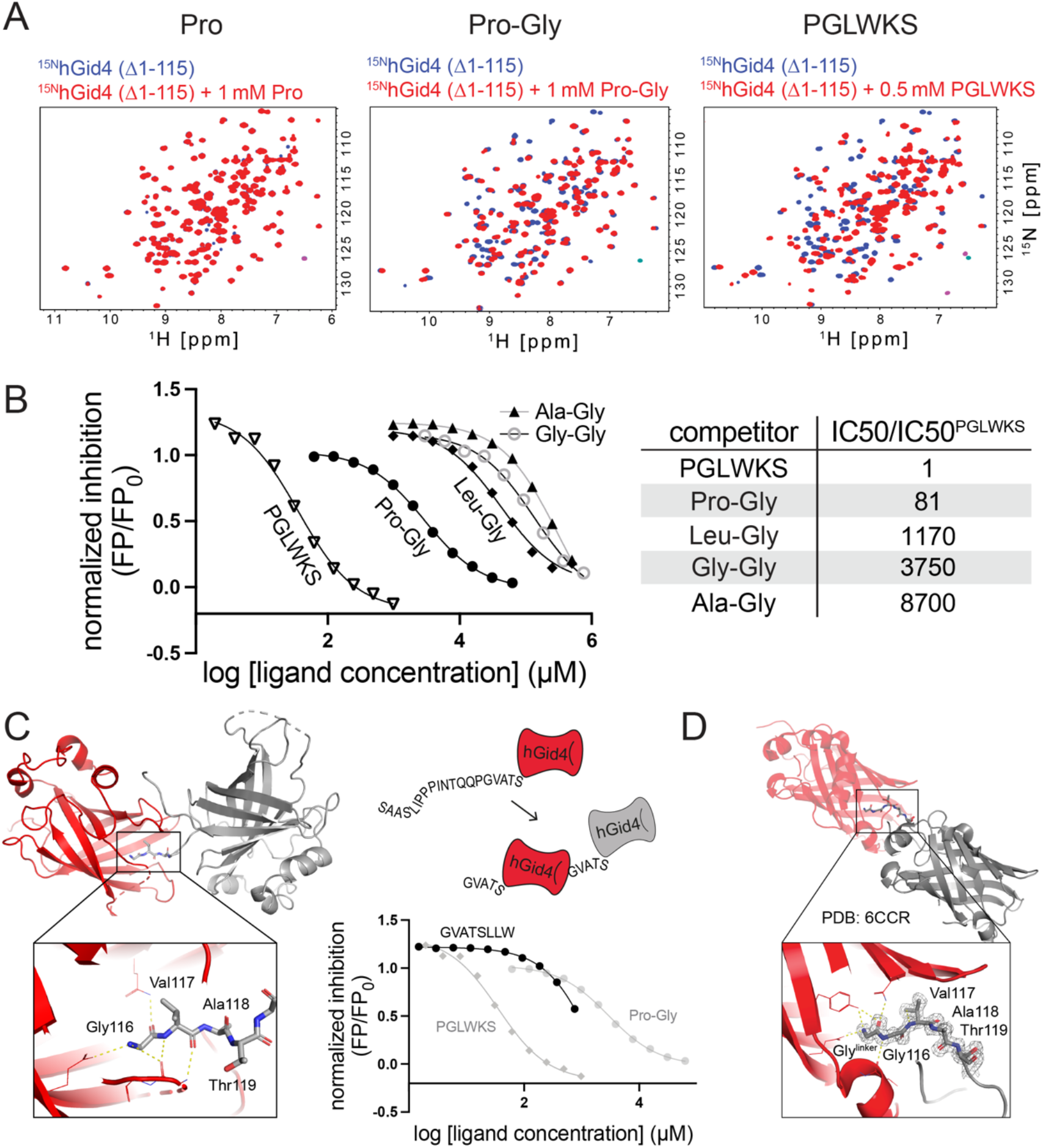
hGid4 recognizes various peptide N-termini and several downstream residues. A. Overlaid ^1^H, ^15^N-HSQC NMR spectra of 0.1 mM [^15^N]-labeled 6xHis-hGid4 (Δ1-115) alone (blue) and upon addition of 1 mM Pro, 1 mM Pro-Gly or 0.5 mM PGLWKS peptide (red). B. Competitive fluorescence polarization (FP) experiments comparing different unlabeled ligands for inhibiting hGid4 (Δ1-115) binding to C-terminally fluorescein-labeled PGLWKS peptide. Ratios of FP signals at varying concentrations of unlabeled ligands to that in the absence of a competitor (FP/FP_0_) were plotted as a function of log_10_[ligand concentration] (left). Half-maximal inhibitory concentrations (IC50) for each ligand were determined by fitting to log[inhibitor] vs. response model and presented relative to IC50 of unlabeled PGLWKS peptide (right). C. Crystal structure of one hGid4 (red) accommodating serendipitously generated Gly116-initiating N-terminus of an adjacent hGid4 molecule (grey) in the crystal lattice. The binding strength of the newly generated N-terminal sequence (116-122) to hGid4 was compared to that of PGLWKS and Pro-Gly with competitive FP (right bottom). D. Previously published hGid4 crystal structure (PDB: 6CCR) revealing one hGid4 binding the N-terminus bearing an additional Gly upstream Gly116 derived from cloning of an adjacent hGid4 molecule (grey) in the lattice of a distinct crystal form.

Given the ability of a Pro-Gly dipeptide to bind hGid4, we examined importance of the N-terminal residue by testing commercially-available variants (Leu-Gly, Ala-Gly, and Gly-Gly along with Pro-Gly) for competing with a fluorescently-labeled PGLWKS peptide whose binding to hGid4 can be measured by fluorescence polarization (FP) (Fig. S1B). Although each of the dipeptides yielded sigmoidal curves, those with N-terminal Pro or Leu were superior (Fig. 1B). Pro-Gly showed a 15-fold lower IC_50_ than Leu-Gly, consistent with prior studies emphasizing the importance of an N-terminal Pro [39].

To examine roles of individual positions in the 6-residue PGLWKS sequence, we employed peptide spot arrays testing all natural amino acids in position 1, positions 2 and 3 together, position 4 or position 5 (Fig. S1C). Binding was detected after incubating the membranes with the substrate binding domain of hGid4, and immunoblotting with anti-hGid4 antibodies. Overall, the data confirm the previous findings that out of the peptides tested PGLWKS is an optimal binder, and that N-terminal non-Pro hydrophobic residues are tolerated in the context of the downstream GLWKS sequence albeit with lower binding [10, 39].

The peptide array data also highlighted the importance of context. Amongst the 400 possible combinations of residues 2 and 3, Gly is preferred at position 2 and Ile or Leu at position 3, mirroring the previously defined sequence preferences. The dynamic range of our assay suggested that downstream residues also contribute to specificity, by unveiling pronounced amino acid preference for bulky hydrophobics and some non-hydrophobic residues also at position 4. In agreement with the structural data [10], the 5^th^ position following the PGLW sequence tolerates many amino acids.

Despite this seemingly strong preference for an N-terminal Pro, we serendipitously visualized hGid4 recognizing a supposedly non-cognate sequence when we set out to visualize its structure in the absence of a peptide ligand by X-ray crystallography. Unexpectedly, the electron density from data at 3 Å resolution showed the first visible N-terminal residue of one molecule of hGid4 inserted into the substrate binding tunnel of an adjacent hGid4 molecule in the crystal lattice (Fig. 1C; Table S1). Perplexingly, this was not the first residue of the input hGid4 construct but Gly116 located 16 positions downstream. It appears that hGid4 underwent processing during crystallization, although it remains unknown how this neo-N-terminus was generated. Nonetheless, the potential for hGid4 to recognize a non-cognate N-terminal Gly was supported by re-examination of the published “apo” hGid4 crystal. In 6CCR.PDB, distinct crystal packing is also mediated by a peptide-like sequence (initiating with a Gly from the Tobacco Etch Virus (TEV) protease cleavage site, followed by hGid4 Gly116) inserting into the substrate binding tunnel of the neighboring molecule in the lattice (Fig. 1D). We speculate that these structurally-observed interactions were favored by the high concentration of protein during crystallization.

To test binding of our fortuitously identified hGid4-binding sequence in solution, we examined competition with the fluorescently-labeled PGLWKS peptide (Fig. 1C). Low solubility of the GVATSLLW peptide (hGid4 residues 116-122) precluded accurate measurement of IC_50_ using our competitive FP assay. Nonetheless, the data qualitatively indicated that the GVATSLLW peptide binds to hGid4 with lower affinity than PGLWKS, but more tightly than the Pro-Gly dipeptide.

Taken together with published work, the data confirmed hGid4’s preference for binding to the previously-defined sequence PGLWKS, but they also highlighted capacity for hGid4 to recognize alternative N-termini. Moreover, given that specific combinations of residues downstream of the Pro-Gly substantially impact the interaction, we considered the possibility that hGid4 recognition of N-terminal sequences could be influenced by context.

### Identification of superior hGid4-binding motifs not initiated by Pro

To discover alternative hGid4-binding sequences that do not initiate with Pro, we constructed a highly diverse N-terminal peptide phage-displayed library of 3.5×10^9^ random octapeptides. The library was constructed after the signal peptide using 8 consecutive NNK degenerate codons encoding for all 20 natural amino acids and fused to the N-terminus phage coat protein. It is expected that Arg or Pro located next to the cleavage site (position +1) will be inexistent or strongly underrepresented because they are known to either inhibit the secretion of phages [40, 41] or the signal peptidase cleavage [42, 43], respectively.

The library was cycled through five rounds of selections following an established protocol [44] to enrich for phages displaying peptides that preferentially bound hGid4 (Fig 2A).

**Figure 2:**
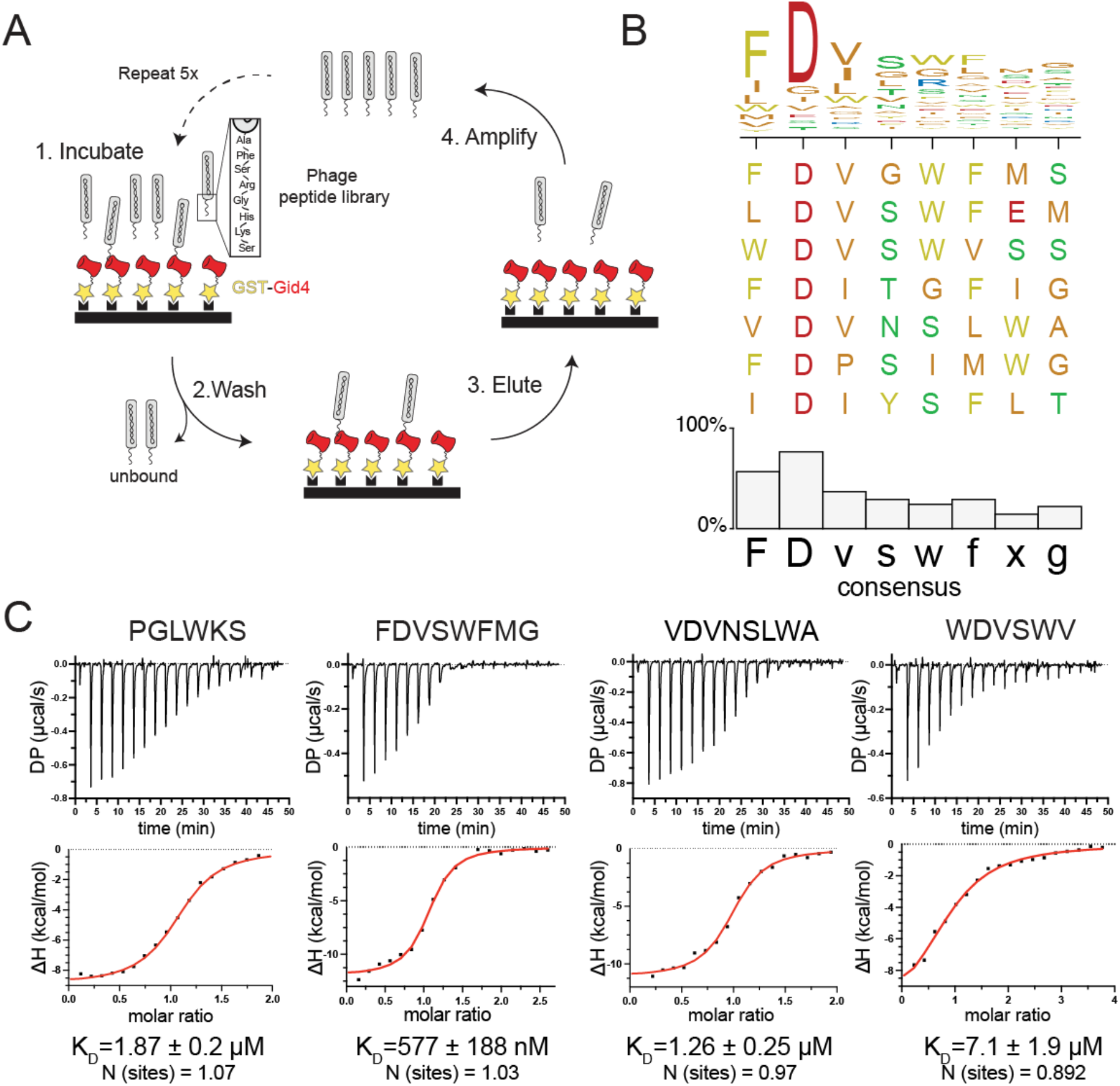
Identification of a high-affinity hGid4-binding motif initiating with non-Pro hydrophobic residues. A. Schematic of phage-display peptide library screen identifying peptides binding GST-tagged hGid4 B. Consensus motif obtained from multiple sequence alignment of 41 unique hGid4-binding peptide sequences (out of which a representative set of 7 sequences is shown). The height of the bars reflects the frequency of a given residue at different positions of the consensus. C. Isothermal titration calorimetry (ITC) to quantify binding of newly determined sequences to hGid4 (Δ1-115). The amount of heat released (ΔH) upon peptide injection was calculated from integrated raw ITC data (top) and plotted as a function of peptide:protein molar ratio (bottom). Dissociation constant (K_D_) and the stoichiometry of the binding event (N) were determined by fitting to the One Set of Sites binding model.

Phages from individual clones that bound to GST-hGid4 but not a control GST based on phage ELISA were subjected to DNA sequence analysis.

The screen yielded 41 unique sequences, none of which were overtly similar to the previously defined hGid4-binding consensus motif PGLWKS (Fig. 2B; Table S2). A new consensus emerged with the following preferences: (1) hydrophobic residues at position 1, with Phe predominating; (2) Asp at position 2; (3) hydrophobic residues at positions 3 and 6, and to a lesser extent at position 5; and (4) small and polar residues at positions 4 and 7. Unlike the PGLWKS sequence wherein the striking selectivity is predominantly for the first four residues, this new consensus extends through the seventh residue.

Although peptides with non-Pro hydrophobic N-termini were previously shown to bind hGid4, the tested sequences bound with one to two orders-of-magnitude lower affinity (K_D_ for IGLWKS 16 μM, VGLWKS 36 μM) than to PGLWKS (K_D_=1.9 μM) (Fig. S2A) [39]. To determine how the newly identified sequences compare, we quantified interactions by isothermal titration calorimetry (ITC). Notably, the peptides of sequences FDVSWFMG and VDVNSLWA showed superior binding (K_D_=0.6 and 1.3 μM, respectively) to the best binder with an N-terminal Pro (Fig. 2C and S2A). Moreover, the affinity for a sequence starting with a Trp (K_D_=7.1 μM for WDVSWV) was superior to the previously identified best binders initiating with a non-Pro hydrophobic residue. Thus, hGid4 is able to accommodate even the bulkiest hydrophobic sidechain at the N-terminus of an interacting peptide. Taken together, the data show hGid4 binds a wide range of peptide sequences, with affinity strongly influenced by residues downstream of the N-terminus.

### hGid4 structural pliability enables recognition of various N-terminal sequences

To understand how hGid4 recognizes diverse sequences, we determined its crystal structure bound to the FDVSWFMG peptide (Fig. 3A, Table S1; all peptide residues except C-terminal Gly visible in density). Overlaying this structure with published coordinates for other hGid4 complexes revealed diverse N-termini protruding into a common central substrate-binding tunnel (Fig. S2B, Phe (our study), or Pro, Leu, Val, or newly recognized Gly [10, 39]). The N-terminal amine groups are anchored through contacts with hGid4 Glu237 and Tyr258 at the tip of the substrate binding tunnel, and common hydrogen bonds to the peptide backbone.

**Figure 3:**
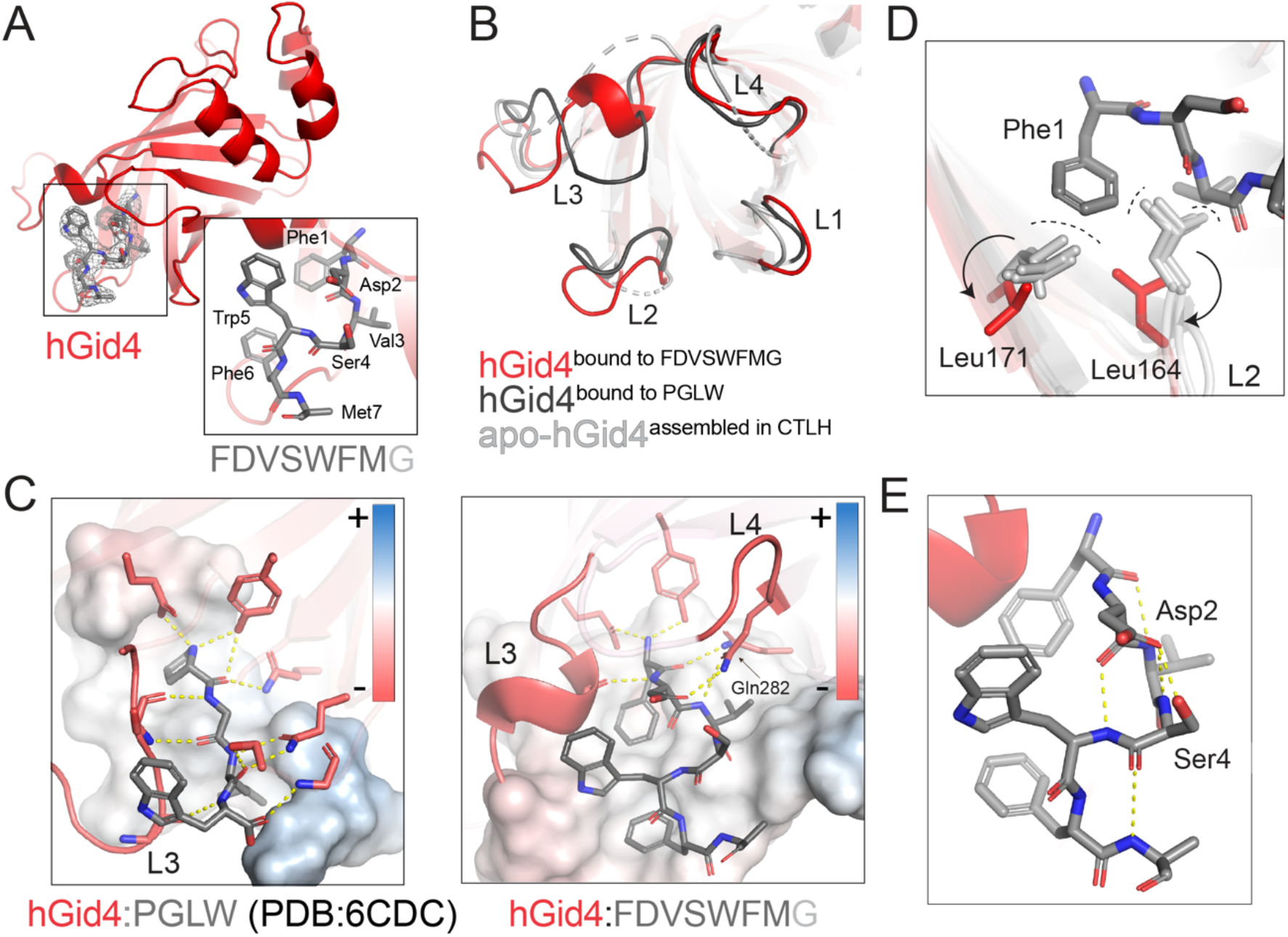
Molecular details of high-affinity peptide binding by hGid4. A. Crystal structure of hGid4 (Δ1-120, Δ294-300) bound to the FDVSWFMG peptide. Clear electron density (2F_O_-F_C_, contoured at 1.5 σ; grey mesh) was visible for all peptide residues besides the C-terminal Gly and the sidechain of Met7, presumably reflecting their mobility. B. Conformations of binding tunnel hairpin loops in apo-hGid4 assembled in CTLH^SR4^ (PDB: 7NSC, light grey) as well as PGLW- (PDB: 6CDC, dark grey) and FDVSWFMG-bound (red) hGid4. C. Comparison of PGLW (left) and FDVSWFMG (right) binding modes to hGid4. Hydrogen bonds between hGid4 residues (red sticks) and peptides (dark grey sticks) are depicted as yellow dashes, whereas the predominantly hydrophobic character of the binding tunnel is visualized as electrostatic potential surface (plotted at ± 7 kT/e; surface colored according to the potential: red – negative (-), blue – positive (+), white – uncharged). D. Overlay of hGid4 bound to PGLW (light grey, PDB: 6CDC), IGLWKS (light grey, PDB: 6WZX), VGLWKS (light grey, PDB: 6WZZ) and FDVSWFMG (red) revealing conformational changes of L2 loop, which prevents steric clash (black dashes) between Leu164 and Leu171 and N-terminal Phe. E. Intrapeptide hydrogen bonding pattern (yellow dashes) within FDVSWFMG upon binding to hGid4.

The structures suggest that the varying peptide sequences are accommodated by complementary conformations of four hairpin loops (L1-L4) at the edge of the hGid4 substrate-binding tunnel (Fig. 3B). The L2, L3, and L4 loops are fully or partially invisible, and are presumably mobile, in the structure of apo-hGid4 assembled in a subcomplex with its interacting subunits from the CTLH E3 [38]. However, they are ordered and adopt different conformations when bound to the different peptides.

As compared to the structure with PGLWKS, the interactions with FDVSWFMG are relatively more dominated by hydrophobic rather than electrostatic contacts (Fig. 3C). The L2 and L3 loops are relatively further from the central axis of the hGid4 β-barrel to interact with more residues in the peptide sequence. The different position of the L2 loop is also required to accommodate the bulkier N-terminal Phe side-chain, compared to Pro or other residues (Fig. 3D). Meanwhile, repositioning of the L4 loop places hGid4 Gln282 to form a hydrogen bond with Asp at the peptide position 2. Moreover, upon binding to hGid4, FDVSWFMG itself adopts a structured conformation owing to multiple intrapeptide backbone hydrogen bonds as well as interaction of Asp2 sidechain with the sidechain and backbone amide of Ser4 (Fig. 3E). Overall, the structures reveal pliability of the hGid4 substrate-binding tunnel enabling interactions with a range of N-terminal sequences, which themselves may also contribute interactions by conformational complementarity.

### Yeast GID substrate receptors recognize natural degrons with suboptimal affinity

To extend our findings to the yeast GID system, we screened the phage peptide library for binders to the yGid4 and yGid10 substrate receptors. The selected consensus sequence binding yGid4 paralleled that for hGid4 (Fig. 4A; Table S2), in agreement with their being true orthologs. Remarkably, despite high similarity to the Gid4s, and its only known endogenous substrate likewise initiating with a Pro [36], the selections with yGid10 identified 12 unique sequences, some with bulky hydrophobic residues and others with Gly prevalent at position 1, each followed by a distinct downstream pattern (Fig. 4B; Table S3). By solving an X-ray structure of yGid10 bound to FWLPANLW peptide and superimposing it on its prior structure with N-terminus of its *bona fide* substrate Art2 [36], we confirmed that the novel sequence is accommodated by the previously characterized binding pocket of yGid10 (Fig. 4C and S3A; Table S1). Moreover, conformations of the yGid10 loops varied in complexes with different peptides, suggesting like hGid4, yGid10 structural pliability allows recognition of various N-terminal sequences (Fig. S3B).

**Figure 4:**
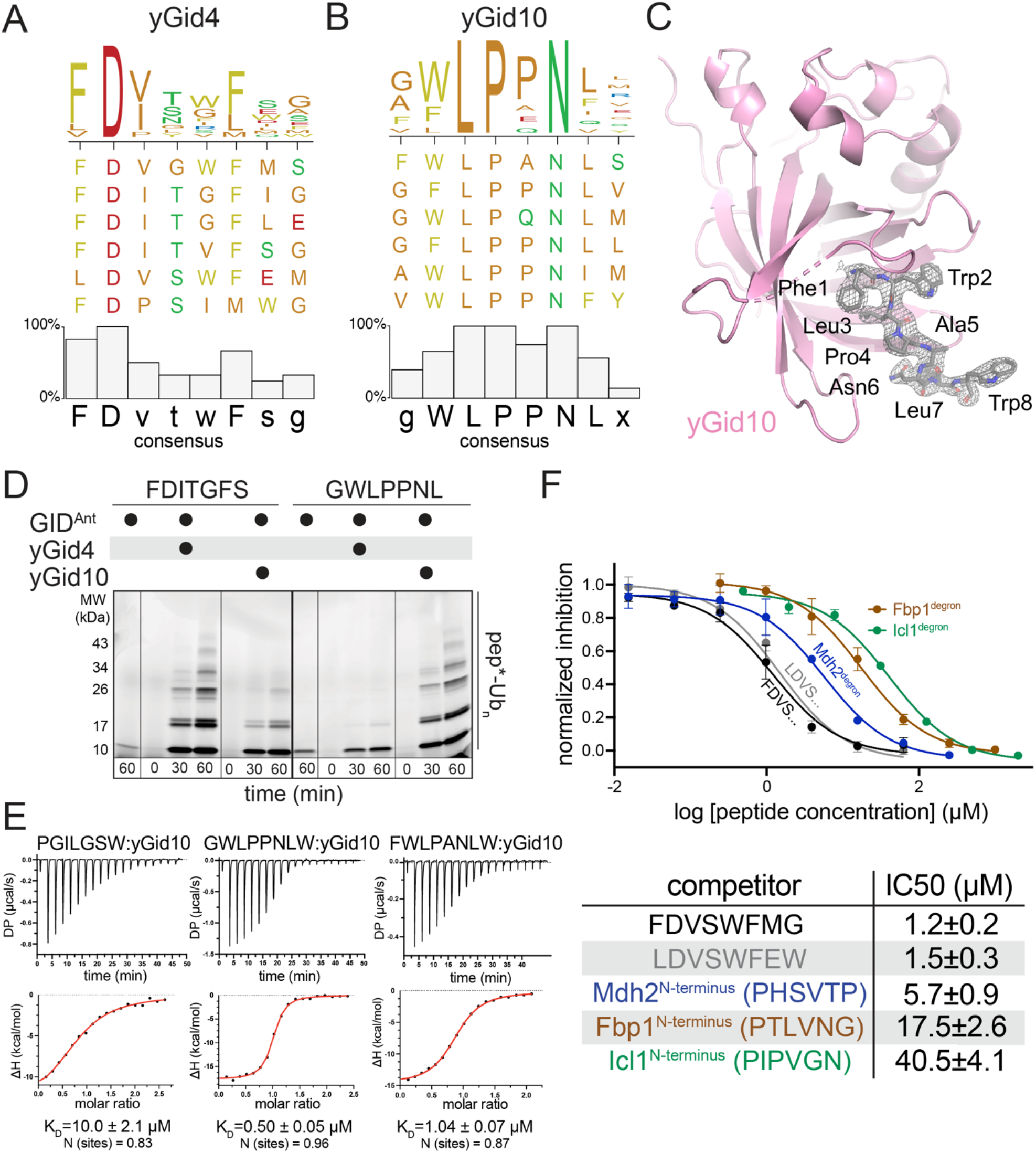
Identification of novel yGid4 and yGid10-binding sequence motifs superior to natural degrons. A. Consensus motif obtained by multiple sequence alignment of 12 unique yGid4-binding peptide sequences B. Consensus motif obtained by multiple sequence alignment of 12 unique yGid10-binding peptide sequences. C. Crystal structure of yGid10 (Δ1-64, Δ285-292) (pink) bound to FWLPANLW (grey sticks). The 2F_O_-F_C_ electron density map corresponding to the peptide is shown as grey mesh contoured at 2σ. D. Fluorescent scans of SDS-PAGE gels after *in vitro* ubiquitylation of fluorescent model peptides harboring either a yGid4 or yGid10-binding sequence by GID^Ant^ (comprising 2 copies each of Gid1 and Gid8, and one copy each of Gid5, Gid2 and Gid9) mixed with either yGid4 (Δ1-115) or yGid10 (Δ1-56) (forming GID^SR4^ or GID^SR10^, respectively). The model peptides contained a corresponding phage display-determined consensus at the N-terminus connected to C-terminal fluorescein (indicated by an asterisk) with a flexible linker. E. ITC binding assays as in Fig. 2C but quantifying binding of several peptides to yGid10 (Δ1-56). F. Competitive *in vitro* ubiquitylation assays probing binding of two novel Phe- and Leu-initiating sequences to yGid4 (Δ1-115) as compared to N-termini of natural GID substrates (Mdh2, Fbp1 and Icl1). Unlabeled peptides were titrated to compete off binding of fluorescent Mdh2 (labeled with C-terminal fluorescein) to GID^SR4^, thus attenuating its ubiquitylation. Normalized inhibition (fraction of ubiquitylated Mdh2 at varying concentration of unlabeled peptides divided by that in the absence of an inhibitor) was plotted against peptide concentration. Fitting to [inhibitor] vs. response model yielded IC50 values and its standard error based on 2 independent measurements.

To test if the selected sequences can mediate binding of substrates for ubiquitylation, we connected a yGid4- and a yGid10-binding sequence to a lysine via a flexible linker designed based on prior structural modeling [38]. The peptides also had a C-terminal fluorescein for detection. Incubating the peptides with either GID^SR4^ or GID^SR10^ and ubiquitylation assay mixes revealed that each serves as a substrate only for its cognate E3 (Fig. 4D).

Finally, we sought to quantitatively compare binding of the new sequences to respective substrate receptors. Affinities of yGid10 for Phe and Gly-initiating sequences, measured by ITC, were, respectively, comparable to and 2-fold greater than for a peptide corresponding to the N-degron of a natural substrate Art2 [36] (Fig. 4E and S3C). Notably, the endogenous degron, and selected sequences, bind yGid10 10- to 20-fold more tightly than the Pro-initiating sequence previously identified by a yeast two-hybrid screen [35]. Although yGid4 is not amenable to biophysical characterization, we could rank-order peptides by inhibition of ubiquitylation of a natural GID^SR4^ substrate Mdh2 (Fig. 4F). Comparing IC_50_ values for the different peptides led to two major conclusions: (1) the phage display-selected sequences are better competitors than N-terminal sequences of endogenous gluconeogenic substrates, and (2) natural substrate N-terminal sequences themselves exhibit varying suppressive effects, with degron of Mdh2 being the most potent, followed by those of Fbp1 and Icl1.

### GID E3 supramolecular assembly differentially impacts catalytic efficiency toward different substrates

We were surprised by the differences in IC_50_ values for the naturally occurring degrons from the best-characterized GID E3 substrates, Fbp1 and Mdh2. We thus sought to compare ubiquitylation of the two substrates, which not only display different degrons but also distinct catalytic domains with unique constellations of lysines. Previous studies showed that ubiquitylation of both substrates depends on coordination of degron binding by yGid4 with placement of specific lysines in the ubiquitylation active site [34, 38]. However, while GID^SR4^ is competent for Mdh2 degradation *in vivo*, a distinct E3 assembly – wherein the Gid7 subunit drives two GID^SR4^ complexes into an oval arrangement (Chelator-GID^SR4^) is specifically required for optimal ubiquitylation and degradation of Fbp1 [38]. Two Gid4 subunits in Chelator-GID^SR4^ simultaneously bind degrons from the oligomeric Fbp1, for simultaneous ubiquitylation of specific lysines on two Fbp1 protomers.

Much like for Fbp1, addition of Gid7 to GID^SR4^ was shown to affect Mdh2 ubiquitylation *in vitro*, albeit in a more nuanced way [38]. As a qualitative test for avid binding to two degrons from Mdh2 (whose dimeric state was confirmed by SEC-MALS (Fig. S4A) and homology modeling (Fig. S4B)) we performed competition assays with monovalent (GID^SR4^ alone or with addition of a truncated version of Gid7 that does not support supramolecular assembly) and bivalent (GID^SR4^ with Gid7 to form Chelator-GID^SR4^) versions of the E3, and lysineless monodentate (Mdh2 degron peptide) and bidentate (Mdh2 dimer) inhibitors (Fig. S4C). While the two inhibitors attenuated ubiquitylation of Mdh2 to a similar extent in reactions with the monovalent E3s, only the full-length Mdh2 complex substantially inhibited the bivalent Chelator-GID^SR4^. This suggested that Chelator-GID^SR4^ is capable of avidly binding to Mdh2.

Thus, we quantified roles of the Fbp1 and Mdh2 degrons by measuring kinetic parameters upon titrating the two different GID E3 assemblies. In reactions with monovalent GID^SR4^, the *K*_m_ for Mdh2 was roughly 3-fold lower than for Fbp1, in accordance with differences in degron binding (Fig. 5A and 5B). Although the higher-order Chelator-GID^SR4^ assembly improved the *K*_m_ values for Fbp1 and for Mdh2, the extents differ such that the values are similar for both substrates. Formation of the higher-order Chelator-GID^SR4^ assembly also dramatically increased the reaction turnover number (*k*_cat_) for Fbp1, with a marginal increase for Mdh2 (8- vs. 1.4-times higher *k*_cat_, respectively), which was already relatively high in the reaction with monomeric GID^SR4^ (Fig. 5C and S4D). Combined with its effects on *K*_m_, formation of the Chelator-GID^SR4^ assembly increased catalytic efficiency (*k*_cat_/*K*_m_) more than 100-times for Fbp1 and only 6-fold for Mdh2, which may rationalize Gid7-dependency of Fbp1 degradation.

**Figure 5:**
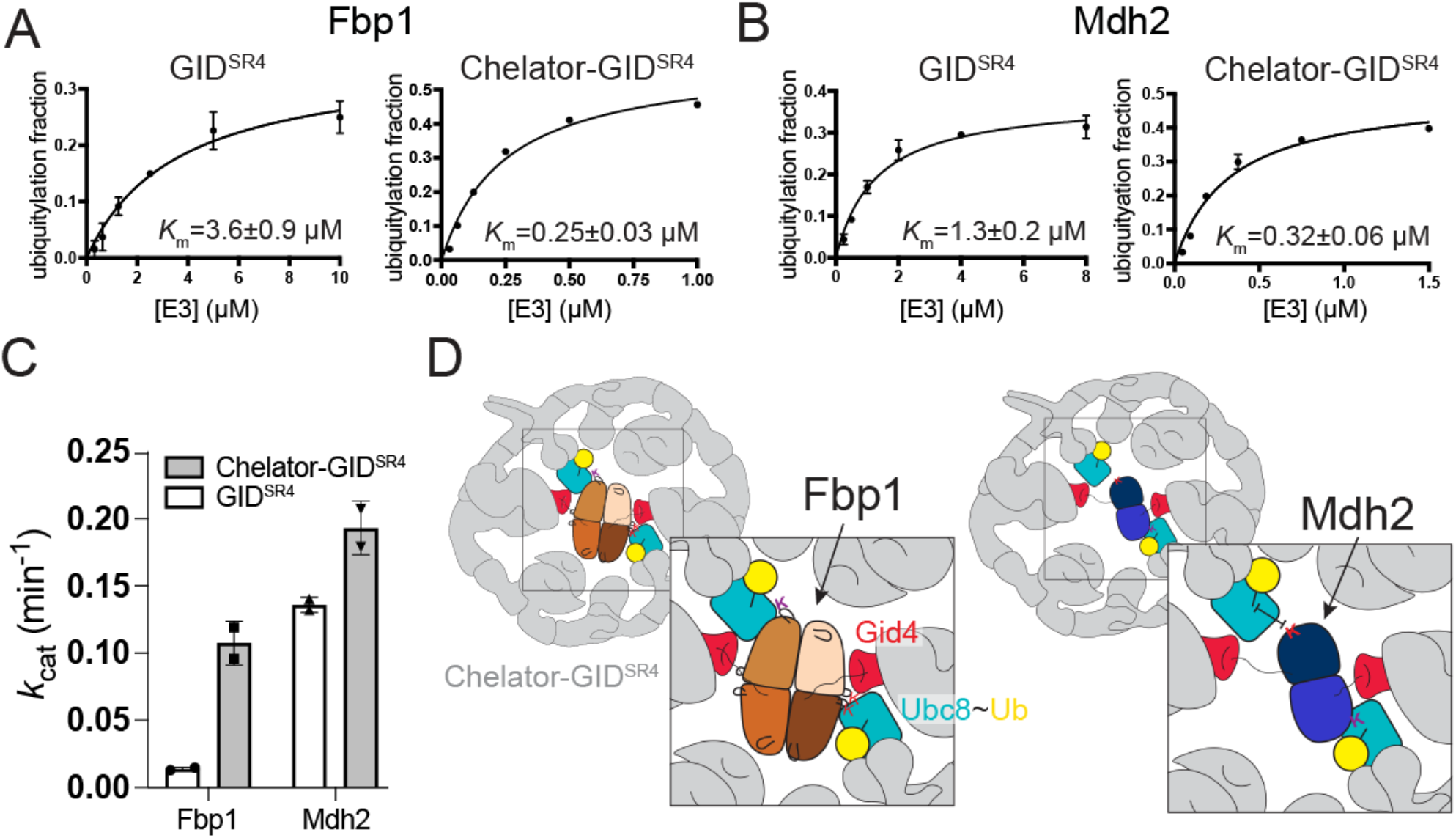
Differential targeting of Mdh2 and Fbp1 by GID E3. A. Plots showing fraction of *in vitro*-ubiquitylated Fbp1 as a function of varying concentration of GID E3 in either its monomeric GID^SR4^ or higher-order Chelator-GID^SR4^ form (co-expressed GID^SR4^ + Gid7). Fitting to Michaelis-Menten equation yielded *K*_m_ values. Error bars represent standard deviation (n=2). B. Plots as in (A) but analyzing Mdh2 ubiquitylation. C. Comparison of *k*_cat_ values for Fbp1 and Mdh2 ubiquitin targeting by GID^SR4^ and Chelator-GID^SR4^ based on a time-course of substrate ubiquitylation (Fig. S4D). D. Cartoons representing ubiquitylation of Fbp1 and Mdh2 by Chelator-GID^SR4^ based on structural modeling (Fig. S5B and S5C).

Beyond avid substrate binding, the multipronged targeting of Fbp1 by Chelator-GID^SR4^ involves proper orientation of the substrate so that specific lysines in metabolic regulatory regions are simultaneously ubiquitylated [38]. To explain the lesser effect of Chelator-GID^SR4^ on catalytic efficiency toward Mdh2, we examined structural models. Briefly, after docking two substrate degrons into opposing Gid4 protomers, we rotated the tethered substrate to place the targeted lysines in the ubiquitylation active sites (Fig. S5A, S5B and S5C). As shown previously, docking either Fbp1 targeted lysine cluster (K32/K35 and K280/K281) places the other in the opposing active site (Fig. 5D and S5B). For Mdh2, however, although the K360/K361 lysine clusters from both Mdh2 protomers could be modeled as simultaneously undergoing ubiquitylation, the two major targeted lysine clusters (K254/K256/K259 and K330) cannot be simultaneously situated in both Chelator-GID^SR4^ active sites (Fig. 5D, S5A and S5C). Thus, the distinct constellations of targeted lysines may also contribute to differences in efficiency of ubiquitylation.

### Degron identity determines *K*_m_ for ubiquitylation but differentially impacts glucose-induced degradation of Mdh2 and Fbp1

To assess the roles of differential degron binding in the distinct contexts provided by the Fbp1 and Mdh2 experiments, we examined the effects of swapping their degrons. We first performed qualitative ubiquitylation assays using the simpler GID^SR4^ E3 ligase. Comparing ubiquitylation of fluorescently-labeled Fbp1 and Mdh2 side-by-side showed more Mdh2 is ubiquitylated with more ubiquitins during the time-course of reactions [38]. These properties are reversed when the N-terminal sequence of Mdh2 is substituted for the Fbp1 degron and vice-versa (Fig. 6A).

**Figure 6:**
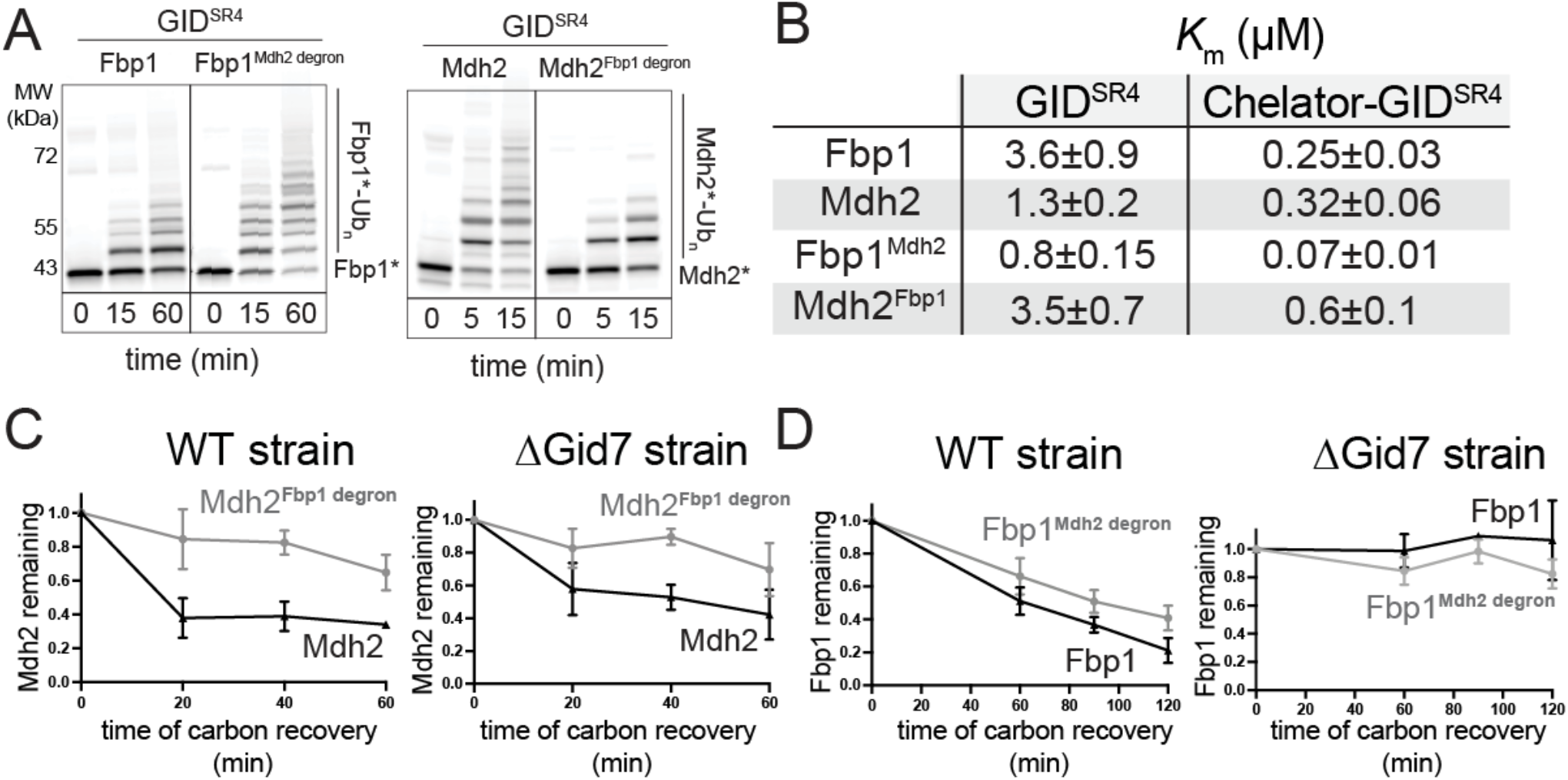
Combinatorial nature of substrate recognition by GID. A. Qualitative *in vitro* ubiquitylation assay probing effect of degron exchange between Fbp1 and Mdh2. Both WT and degron-swapped versions of Fbp1 and Mdh2 were C-terminally labelled with fluorescein (indicated by an asterisk) and ubiquitylated by GID^SR4^. B. Table summarizing values of *K*_m_ for ubiquitylation of WT and degron-swapped substrates by the two versions of GID. C. *In vivo* glucose-induced degradation of exogenously expressed and C-terminally 3xFlag-tagged Mdh2 as well as its degron-swapped versions quantified with a promoter-reference technique. Levels of the substrates (relative to the level of DHFR) at different timepoints after switch from gluconeogenic to glycolytic conditions were divided by their levels before the switch (timepoint 0). For each substrate, the experiment was performed in WT and ΔGid7 yeast strains. Error bars represent standard deviation (n=3), whereas points represent the mean. D. *In vivo* assay as in (C) but with WT and degron-swapped Fbp1.

Quantifying the *K*_m_ values showed that the values for degron-swapped substrates roughly scaled with degron identity (Fig. 6B and S4E; for Mdh2 *K*_m_≈1.3 μM, for degron-swapped Fbp1^Mdh2 degron^ *K*_m_≈0.8 μM, for Fbp1 *K*_m_≈3.6 μM, for degron-swapped Mdh2^Fbp1 degron^≈3.5 μM). Furthermore, as expected, the *K*_m_ values for all substrates improved in reactions with Chelator-GID^SR4^. However, the relative impact seemed to scale with the way in which they are presented from the folded domain of a substrate rather than the degrons themselves (roughly 14-fold for Fbp1 and 11-fold for Fbp1^Mdh2 degron^ versus 4-fold for Mdh2 and 6-fold for Mdh2^Fbp1 degron^).

Effects *in vivo* were examined by monitoring glucose-induced degradation of the wild-type and mutant substrates. Degradation was examined using the promoter reference technique, which normalizes for translation of an exogenously expressed substrate (here, C-terminally 3xFLAG-tagged versions of Fbp1, Mdh2, Fbp1^Mdh2 degron^ and Mdh2^Fbp1 degron^) relative to a simultaneously expressed control [7, 45]. As shown previously, Mdh2 was rapidly degraded in the wild-type yeast and the ΔGid7 strain (Fig. 6C) [38]. However, turnover of the mutant version bearing the weaker Fbp1 degron was significantly slower in both genetic backgrounds. Thus, the Mdh2 degron is tailored to the Mdh2 substrate. In striking contrast, although the Mdh2 degron did subtly impact degradation of Fbp1, it was not sufficient to overcome dependency on Gid7 (Fig. 6D). Thus, substrate ubiquitylation, and turnover, depend not only on degron identity, but also on their associated targeted domains.

## DISCUSSION

Overall, our study of degron recognition by the Gid4/Gid10-family of substrate receptors of GID/CTLH E3s leads to several conclusions. First, GID/CTLH E3 substrate receptors recognize a diverse range of N-terminal sequences, dictated not only by the N-terminal residue, but also the pattern of downstream amino acids (Fig. 1 and S1). Second, such diverse N-terminal sequence recognition is achieved by the combination of (1) a deep substrate-binding tunnel culminating in a conserved Glu and Tyr placed to recognize the N-terminal amine (2) pliability of loops at the entrance to the substrate binding tunnel that can conform to a range of downstream sequences, and (3) formation of distinct extended structures by the N-terminal peptide sequences that can complement the receptor structures (Fig. 3). Remarkably, the hGid4 loops and the bound peptide reciprocally affect each other – peptide binding induces folding of the flexible loops in a conformation that depends on a peptide sequence, whereas the arrangement of the loops dictates affinity for the bound peptide by altering shape and properties of the binding pocket. This correlation rationalizes strong dependence of Gid4 specificity on the peptide sequence context. For instance, in the conformation induced by XGLWKS, hGid4 is strikingly specific towards N-terminal Pro, whereas the novel XDVSWFMG sequence opens up the binding pocket and favors bulky hydrophobics at position 1. Third, the range of interactions result in a range of affinities (Fig. 2, 4 and S2A). Notably, our randomized phage-display peptide library screen identified far tighter binders to yGid4 than known natural degrons, which themselves bind yGid4 with varying K_D_s. This approach identified yGid10-binding sequences on par with the only known natural degron, and with higher affinity than a sequence identified by yeast two-hybrid screening. Phage-display peptide library screening may thus prove to be a useful method for degron identification. Fourth, degron binding is only part of substrate recognition by GID E3s (Fig. 5 and 6). Rather, ubiquitylation and degradation depend on both the pairing of a degron with a substrate domain that presents lysines in a particular constellation, and configuration of the GID E3 in either a simplistic monovalent format or in a multivalent chelator assembly for optimal targeting.

Some features of the high-affinity peptide binding by Gid4s and yGid10 parallel other end-degron E3s. In particular, several cullin-RING ligases, the founding family of multiprotein E3s with interchangeable substrate receptors [46], have recently been discovered to recognize either specific N- or C-terminal sequences as degrons [9, 47–50]. Structures showed half a dozen or more residues in C-degron sequences engaging deep clefts or tunnels in their cullin-RING ligase substrate receptors, thus conceptually paralleling the high-affinity interactions with Gid4s and yGid10 [11, 51–54]. Furthermore, much like Gid4s and Gid10 recognize diverse sequences, the substrate-binding site of a single cullin-RING ligase was recently shown to bind interchangeably to a C-degron or to a different substrate’s internal sequence [53–55]. Another interesting parallel between Gid4/Gid10-type recognition and some Ubr-family E3s is potential to bind diverse N-terminal sequences. However, while our data show that a common binding cleft in the Gid4/Gid10 fold structurally accommodates diverse sequences with high-affinity, some Ubr-family E3s bind different N-terminal sequences using distinct N-degron-binding domains [56–58]. Moreover, many structurally-characterized N-terminal peptide-bound E3s have shown shallower modes of recognition. Structures including the UBR-box 1 [59–62] and UBR-box2-type recognition by bacterial [63–65] and plant [66] ClpS homologs revealed this strict degron specificity is determined by as few as two or three amino acids.

To-date, few GID E3 substrates have unambiguously been identified. Thus, our findings have implications for identification of new substrates. Most of the currently characterized substrates depend on co-translational generation of an N-terminal Pro. However, sequences initiating with bulky hydrophobic residues may be refractory to N-terminal processing enzymes such as Met aminopeptidase [67–69]. Nonetheless, post-translational processing could generate such N-termini. Several paradigms for post-translational generation of N-degrons have been established by studies of Ubr1 substrates. First, endoproteolytic cleavage – by caspases, calpains, separases, cathepsins and mitochondrial proteases [37, 70–75] - is responsible for the generation of myriad Arg/N-degron pathway substrates recognized by some Ubr-family E3s [2]. Notably, over 1800 human proteins have an FDI/V sequence within them, raising the possibility that the newly identified Gid4-and Gid10-binding motifs likewise could be exposed upon post-translational protein cleavage. Second, some N-degrons are created by aminoacyl-tRNA protein transferases-catalyzed appendage of an additional amino acid at the protein’s N-terminus [58, 76–79]. In higher eukaryotes, UBR1 and UBR2 associating with such post-translational N-terminal arginylation and other enzymes in a supercomplex was proposed to enable substrate superchanelling between the enzyme generating the degron and that recognizing it [80]. The bacterial N-degron pathway involves conjugation of hydrophobic residues such as Phe and Leu [81–85]. It is tempting to speculate that hydrophobic N-degrons in eukaryotes could involve N-terminal amino acid addition. Finally, yeast Ubr1 is modulated in an intricate manner: after HtrA-type protease cleavage, a portion of the protein Roq1 binds Ubr1 and alters its substrate specificity [86]. Notably, proteomic studies showed that the human CTLH complex itself associates with the HtrA-type protease HTRA2 [22, 24, 87–89], known to be involved in mitochondrial quality control [90, 91]. This raises the tantalizing possibility that the CTLH E3 might form a multienzyme targeting complex that integrates a regulatory cascade to generate its own substrates or regulatory partners.

Finally, our examination of degron-swapped actual GID E3 substrates Fbp1 and Mdh2 showed that N-terminal sequence is only part of the equation determining ubiquitylation and subsequent degradation. Mdh2 required its own degron and its ubiquitylation and degradation were impaired when substituted with the weaker degron from Fbp1, irrespective of capacity for GID^SR4^ to undergo Gid7-mediated superassembly. However, while either degron could support Fbp1 targeting, this requires Gid7-dependent formation of the Chelator-GID^SR4^ supramolecular chelate-like E3 configuration. Taken together, our data reveal that structural malleability of both the substrate receptor and the E3 supramolecular assembly endows GID E3 complexes – and presumably CTLH E3s as well – capacity to conform to diverse substrates, with varying degrons and associated targeted domains. Such structural malleability raises potential for regulation through modifications or interactions impacting the potential conformations of both the substrate binding domains and higher-order assemblies, and portends future studies will reveal how these features underlie biological functions of GID/CTLH E3s across eukaryotes. Moreover, our results highlight that turnover depends on structural complementarity between E3 and both the substrate degron and ubiquitylated domains, a principle of emerging importance for therapeutic development of targeted protein degradation.

## ACKNOWLDEGMENTS

We thank Chen G. and Pavlenco A. for construction of N-terminal peptide phage-displayed library; A. Varshavsky for promoter reference plasmids; S. Uebel and S. Pettera for peptide synthesis; K. Valer-Saldana and S. Pleyer for assistance with protein crystallization; J. Rech for the preparation of peptide spot arrays; Paul Scherrer Institut, Villigen, Switzerland for provision of synchrotron radiation beamtime at beamlines PXII and X10SA of the SLS; G. Kleiger for guidance regarding kinetics; I. Paron for technical assistance with mass spectrometry; S.v. Gronau for maintenance of insect cells and the Schulman lab for advice and support.

The project was funded by the Deutsche Forschungsgemeinschaft (DFG) SFB1035 (B.A.S. and M.S.), and Leibniz Prize SCHU 3196/1 (B.A.S.). Work in the Schulman lab is supported by the Max Planck Society.

## AUTHOR CONTRIBUTIONS

Initial conceptualization: J.C., D.S., S.S.D. and B.A.S.; methodology: J.C., D.S., J.T., G.P., J.B., M.S., J.R.P., S.S.D. and B.A.S.; investigation: J.C., D.S., J.T., M.S.L., G.P., J.B., and J.R.P.; resources: J.C., D.S., J.T. and M.S.L.; writing: J.C., D.S. and B.A.S.; supervision: M.S., S.S.D. and B.A.S.; funding acquisition: M.S., S.S.D and B.A.S.

## METHODS

### Reagent table

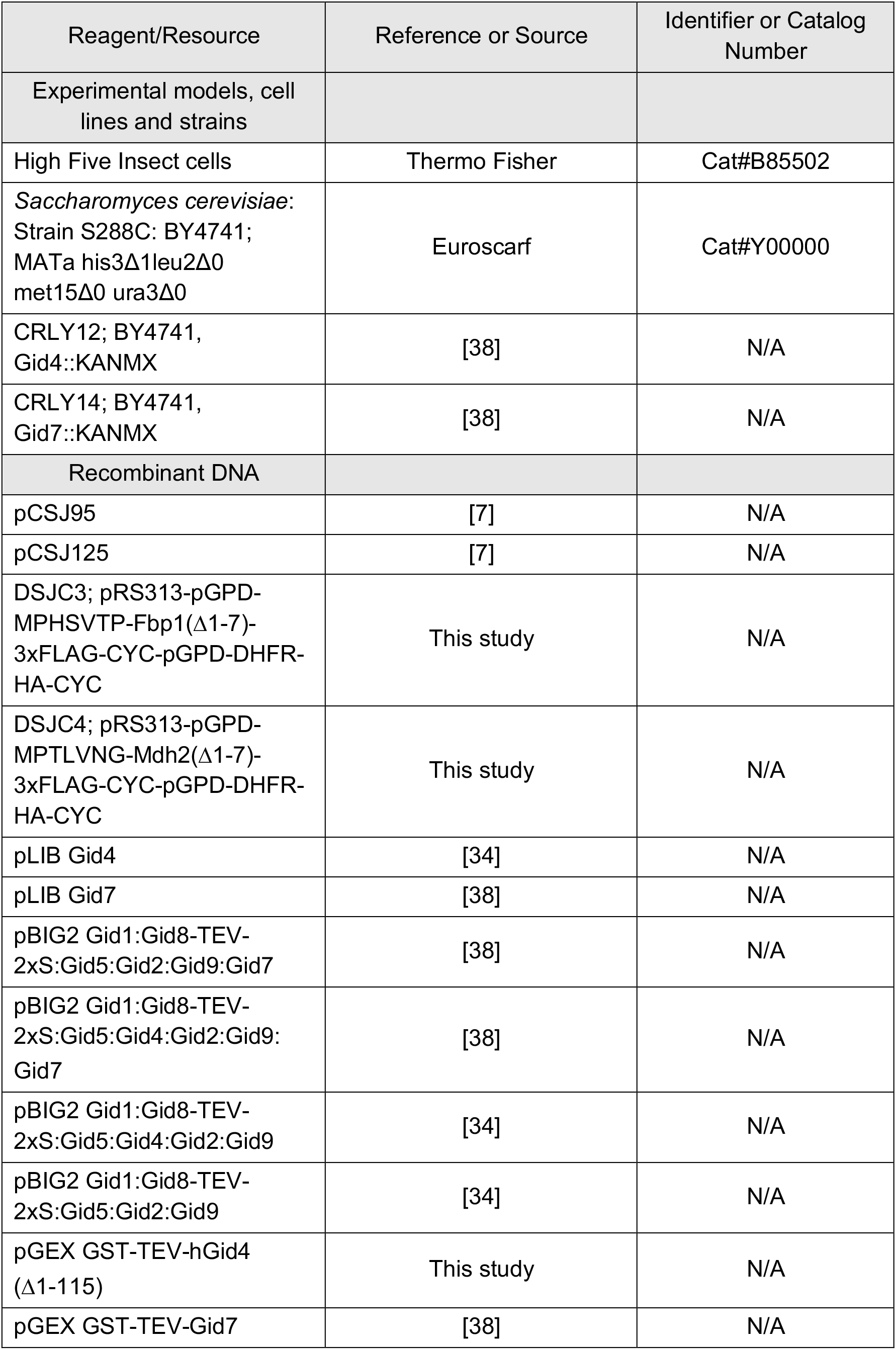
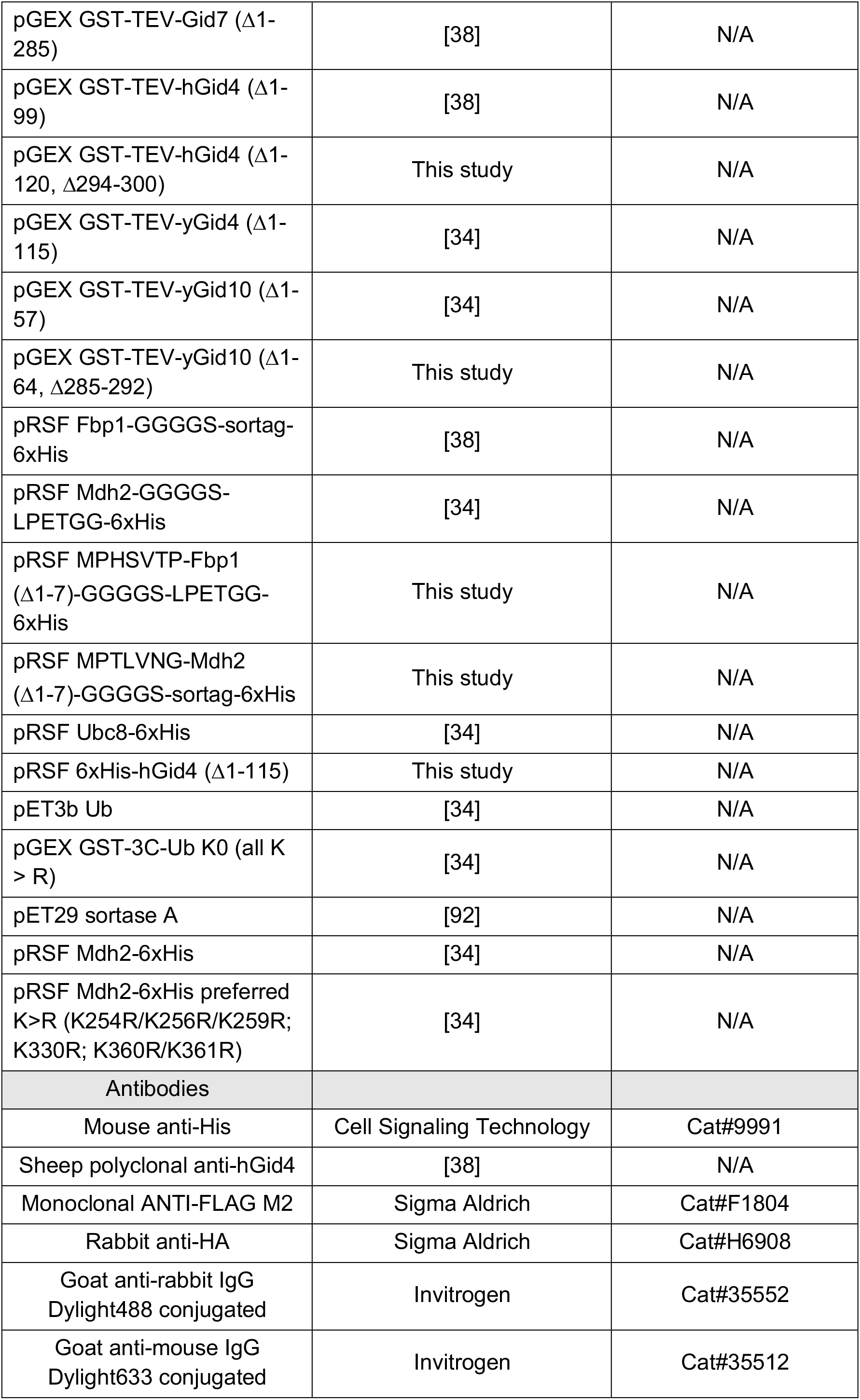
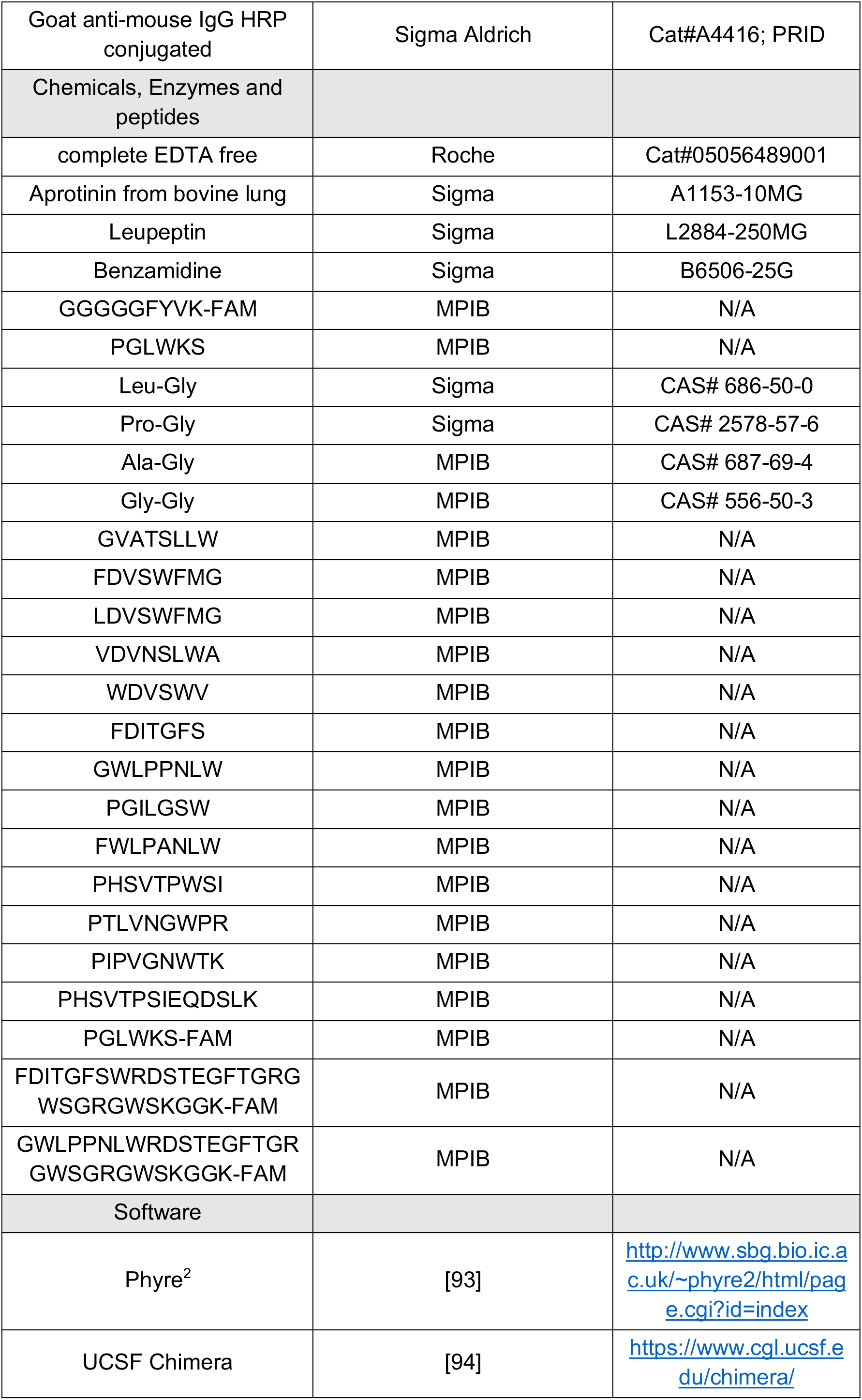
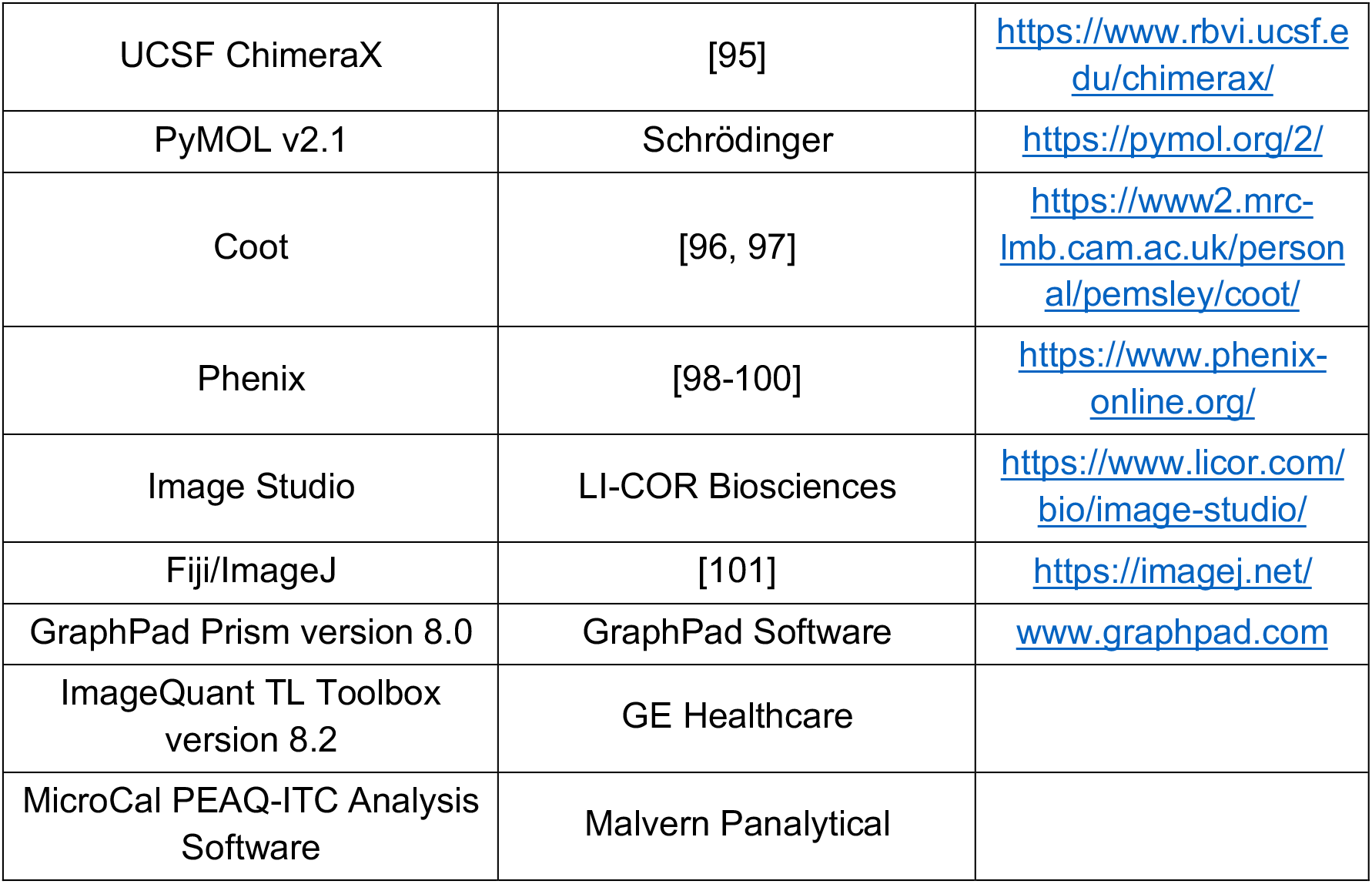

### Data availability

The accession codes for the PDB models will be available in RCSB. All the unprocessed image data will be deposited to Mendeley Data.

### Plasmid preparation and mutagenesis

All the genes encoding yeast GID subunits including the substrate receptors yGid4 and yGid10, as well as Fbp1 and Mdh2 substrates were amplified from *S. cerevisiae* BY4741 genomic DNA. The gene encoding hGid4 was codon-optimized for bacterial expression system and synthesized by GeneArt (Thermo Fisher Scientific).

All the recombinant constructs used for protein expression were generated by Gibson assembly method [102] and verified by DNA sequencing. The GID subunits were combined using the biGBac method [103] into a single baculoviral expression vector. All the plasmids used in this study are listed in the Reagent table.

### Bacterial protein expression and purification

All bacterial expressions were carried out in E. coli BL21 (DE3) RIL cells in a Terrific Broth medium [104] overnight at 18°C. All versions of yGid4, yGid10 and hGid4 (besides that used for NMR) were expressed as GST-TEV fusions. The harvested cell pellets were resuspended in the lysis buffer (50 mM HEPES pH 7.5, 200 mM NaCl, 5 mM DTT and 1 mM PMSF), disintegrated by sonication and subjected to glutathione affinity chromatography, followed by overnight cleavage of the pull-downed proteins at 4°C with tobacco etch virus (TEV) protease to release the GST tag. Final purification was performed with size exclusion chromatography (SEC) in the final buffer containing 50 mM HEPES pH 7.5, 200 mM NaCl and 1 mM or 5 mM DTT (for assays and crystal trials, respectively), or 0.5 mM TCEP (for ITC binding assay). Additionally, pass-back over glutathione affinity resin was performed in order to get rid of the remaining uncleaved GST-fusion protein and free GST.

All versions of Ubc8, Fbp1 and Mdh2 were expressed with a C-terminal 6xHis tag. The harvested cell pellets were resuspended in the lysis buffer (50 mM HEPES pH 7.5, 200 mM NaCl, 5 mM β-mercaptoethanol, 10 mM imidazole and 1 mM PMSF) and sonicated. Proteins were purified by nickel affinity chromatography, followed by anion exchange and SEC in the final buffer containing 50 mM HEPES pH 7.5, 200 mM NaCl and 1 mM DTT.

Untagged WT ubiquitin used for *in vitro* assays was purified via glacial acetic acid method [105], followed by gravity S column ion exchange chromatography and SEC.

### Insect cell protein expression and purification

All yeast GID complexes used in this study were expressed in insect cells. For protein expression, Hi5 insect cells were transfected with recombinant baculovirus variants and grown for 60 to 72 hours in EX-CELL 420 Serum-Free Medium at 27°C. The insect cells were harvested by centrifugation at 450xg for 15 mins and pellets were resuspended in a lysis buffer (50 mM HEPES pH 7.5, 200 mM NaCl, 5 mM DTT, 10 μg/ml leupeptin, 20 μg/ml aprotinin, 2 mM benzamidine, EDTA-free Complete protease inhibitor tablet (Roche, 1 tablet per 50 ml of buffer) and 1 mM PMSF). All the complexes were purified from insect cell lysates by StrepTactin affinity chromatography by pulling on a twin-Strep tag fused to the Gid8 C-terminus. Further purification was performed by anion exchange chromatography and SEC in the final buffer containing 25 mM HEPES pH 7.5, 200 mM NaCl and 1 mM DTT.

### Preparation of fluorescent substrates for *in vitro* activity assays

C-terminal labelling of Fbp1, Mdh2 and their degron-swapped versions with fluorescein was performed through a sortase A-mediated reaction. The reaction mix contained 50 **μ**M substrate (C-terminally tagged with a sortag (LPETGG) followed by a 6xHis tag), 250 **μ**M of a fluorescent peptide (GGGGGFYVK-FAM), 50 **μ**M of sortase A [92] and a reaction buffer (50 mM Tris pH 8, 150 mM NaCl and 10 mM CaCl_2_). The reaction was carried out at room temperature for 30 minutes. After the reaction, a pass-back on Ni-NTA resin was done to get rid of unreacted substrates. Further purification was done with SEC in the final buffer containing 50 mM HEPES pH 7.5, 200 mM NaCl and 1 mM DTT.

### ^15^N labelling of hGid4

For NMR experiments, ^15^N-labeling of 6xHis-hGid4 (Δ1-115) was carried out. Firstly, 50 ml of the preculture was spun at 3000 rpm for 20 mins. The supernatant was then removed and resuspended with 1x M9 cell growth medium (1 g NH_4_Cl, 2 g glucose, 5 mg/ml thiamine chloride, 1 M MgSO_4_, 1 M CaCl_2_ and 1g ^15^NH_4_Cl per liter of 1x M9 medium) containing all essential ions and antibiotics. The cultures were then grown at 37°C and 200 rpm until it reached the OD_600_ of 0.5-0.8. Subsequently, the temperature was reduced to 23°C and kept for an hour before inducing with 0.6 M IPTG. The cultures were then kept growing overnight at 23°C, 200 rpm and harvested the next day following protein purification as described in the section “Protein expression and purification” but in the final SEC buffer containing 25 mM phosphate buffer pH 7.8, 150 mM NaCl and 1 mM DTT.

### NMR (Nuclear Magnetic Resonance) spectroscopy

NMR experiments were recorded at 298 K on Bruker Avance III 600 MHz spectrometer (at ^1^H Larmor frequency of 600 MHz) equipped with a 5 mm TCI cryoprobe. Samples at 0.1 mM ^15^N-labeled hGid4 were prepared in NMR buffer (50 mM HEPES, 100 mM NaCl, pH 7.0) supplemented with 10% D_2_O. ^1^H,^15^N HSQC (heteronuclear single quantum coherence) correlation spectra were acquired with 2048 x 256 complex points and a recycle delay of 1.2 s, with 24 scans. DMSO references were acquired at the beginning and end of the assay. No differences were observed between them. Spectra in the presence of ligands where measured at 1 mM concentration of Pro or Pro-Gly and 0.5 mM of the PGLWKS peptide.

### Phage-displayed N-terminal peptide library construction and selections

A diverse octapeptide N-terminal phage-displayed library was generated for the identification of peptides binding to hGid4 (Δ1-99), yGid4 (Δ1-115) and yGid10 (Δ1-56). An IPTG-inducible P*_tac_* promoter was utilized to drive the expression of open-reading frames encoding the fusion proteins in the following form: the stII secretion signal sequence, followed by a random octapeptide peptide, a GGGSGGG linker and the M13 bacteriophage gene-8 major coat protein (P8). The libraries were constructed by using oligonucleotide-directed mutagenesis with the phagemid pRSTOP4 as the template, as described [106]. The mutagenic oligonucleotides used for library construction were synthesized using with NNK degenerate codons (where N = A/C/G/T & K = G/T) that encode all 20 genetically encoded amino acids. The diversity of the library was 3.5 × 10^9^ unique peptides.

The N-terminal peptide library was cycled through five rounds of binding selections against immobilized GST-tagged hGid4, yGid4, and yGid10, as described [44]. Pre-incubation of the phage pools against immobilized GST was performed before each round of selections to deplete non-specific binding peptides. For rounds four and five, 48 individual clones were isolated and tested for binding to the corresponding targets by phage ELISA [107], and clones with a strong and specific positive ELISA signal were Sanger sequenced. A total of 41, 12, and 12 unique peptide sequences were identified binding to hGid4, yGid4, and yGid10, respectively, and their sequences were aligned to identify common specificity motifs.

Oligonucleotide used for the Kunkel reaction to construct the library: GCTACAAATGCCTATGCANNKNNKNNKNNKNNKNNKNNKNNKGGTGGAGGATCCGGAG GA

### Fluorescence polarization (FP) assays

To determine conditions for a competitive FP assay, we first performed the experiment in a non-competitive format. A 2-fold dilution series of hGid4 (Δ1-115) was prepared in the FP buffer containing 25 mM Tris pH 8.0, 50 mM NaCl, 0.5 mM DTT and 20 nM of fluorescent PGLWKS-FAM peptide. The mixed samples were equilibrated at room temperature for 5 mins before transferring to Greiner 384-well flat bottom black plates. Then, the polarization values were obtained by measuring the excitation at 482 nm and emission at 530 nm using CLARIOstar microplate reader (BMG LABTECH). For each run, the gain was optimized with FP buffer-only control. The data were fit to one site-binding model in GraphPad Prism to determine K_D_ value.

To compare binding of several unlabeled ligands to hGid4, we performed the FP measurements in a competitive format. Based on the FP plot from hGid4 titration experiment, we identified hGid4 concentration, which resulted in ∼60% saturation of the FP signal (6.8 μM hGid4). Next, 2-fold dilution series of unlabeled competitors was prepared in FP buffer mixed with hGid4. After 5 min incubation, the measurement was performed as described above. The data were plotted relative to the FP signal in the absence of an inhibitor as a function of log(ligand concentration) and analyzed with log(inhibitor) vs. response model to determine IC50 values. To determine relative inhibitory strength of the ligands, the determined IC50 values were divided by that of PGLWKS.

### Screening of PGLWKS sequence for hGid4 binding using peptide spot array

The array of peptides derived from the PGLWKS sequence with all 20 amino acid substituted at positions 1, 2 and 3 together, 4 and 5 were synthesized on a membrane in the MPIB biochemistry core facility following the previously established protocols (Hilpert et al., 2007). The membrane blot was first blocked with 3% milk in TBST buffer (20 mM Tris, 150 mM NaCl and 0.1% Tween 20) for 1 hour at room temperature. hGid4 (Δ1-99) was diluted to 10 μg/ml in the buffer containing 150 mM NaCl, 25 mM HEPES pH 7.5, 0.5 mM EDTA pH 8.0, 10% glycerol, 0.1% Tween 20, 2% milk and 1 mM DTT and incubated with the blocked membrane overnight at 4°C with gentle shaking. The membrane was then washed with TBST buffer 3 times, incubated with primary anti-hGid4 sheep monoclonal antibody (1:500) for 3 hours with gentle shaking, followed by multiple washing steps with TBST and 1 hour incubation with secondary HRP-conjugated anti-sheep (1:5000) antibody. The membranes were again washed multiple times with TBST and hGid4 binding was visualized by chemiluminescence in Amersham Imager 800 (Cytiva).

### Isothermal titration calorimetry (ITC) binding assays

To quantify binding of peptides to hGid4 (Δ1-115) and yGid10 (Δ1-56), we employed ITC. All peptides were dissolved in the SEC buffer used for purification of substrate receptors containing 25 mM HEPES pH 7.5, 150 mM NaCl and 0.5 mM TCEP and their concentration was measured by absorbance at 280 nm (a single tryptophan residue was appended at peptides’ C-termini to facilitate determination of peptide concentration). Binding experiments were carried out in the MicroCal PEAQ-ITC instrument (Malvern Pananalytica) at 25°C by titrating peptides to either hGid4 or yGid10. Peptides were added to individual substrate receptors using 19 x 2 µl injections, with 4 s injection time and 150 s equilibration time between the injections. The reference power was set to 10 µcal/s. The concentration of the peptides and substrate receptors were customized according to the ITC plot and the estimated K_D_ values. Raw ITC data were analyzed using One Set of Sites binding model (Malvern Pananalytica) to determine K_D_ and stoichiometry of the binding events (n). All plots presented in figures were prepared in GraphPad Prism.

### Size exclusion chromatography with multiangle light scattering (SEC-MALS)

To determine the oligomeric state of Mdh2, we performed SEC-MALS (conducted in the MPIB Biochemistry Core Facility). For each run, 100 µl of Mdh2 at 1 mg/mL were injected onto Superdex 200 column equilibrated with a buffer containing 25 mM HEPES pH 7.5, 150 mM NaCl and 5 mM DTT.

### *In vitro* activity assays

All ubiquitylation reactions were performed in a multi-turnover format in the buffer containing 25 mM HEPES pH 7.5, 150 mM NaCl, 5 mM ATP and 10 mM MgCl_2_. To quench the reactions at indicated timepoints, an aliquot of the reaction mix was mixed with SDS-PAGE loading buffer. Unless stated otherwise, ubiquitylation of fluorescein-labelled substrates was visualized with a fluorescent scan of an SDS-PAGE gel with a Typhoon imager (GE Healthcare) and quantified with ImageQuant (GE Healthcare; version 8.2).

To verify whether FDITGFS and GWLPPNL can be recognized by, respectively, yGid4 and yGid10 during ubiquitylation reaction (Fig. 4D), we performed an *in vitro* activity assay with model peptides, consisting of the respective N-terminal sequences connected to a single acceptor lysine with a 23-residue linker and C-terminal fluorescein (the length of the linker optimized based on GID^SR4^ structure as in [38]). To start the reaction, 0.2 μM E1 Uba1, 1 μM E2 Ubc8-6xHis, 0.5 μM E3 GID^Ant^, 20 μM Ub, 1 μM yGid4 (Δ1-115) or yGid10 (Δ1-56) and 1 μM peptide substrate were mixed and incubated at room temperature.

In order to probe avid binding of Mdh2 to Chelator-GID^SR4^, we employed a competition ubiquitylation assay (Fig. S4C). The reactions were initiated by mixing 0.2 µM Uba1, 1 µM Ubc8-6xHis, 0.5 µM E3 GID^SR4^, 0 or 2 µM Gid7 (WT or its N terminal deletion mutant, Δ1-284), 0.5 µM of Mdh2-FAM, 20 µM of an unlabeled competitor (dimeric Mdh2-6xHis or a peptide comprising Mdh2 N-terminal sequence PHSVTPSIEQDSLK) and 20 µM Ub. GID^SR4^ was incubated with Gid7 for 5 minutes on ice before the start of the reaction.

In order to test whether the preferred ubiquitylation sites within Mdh2 determined previously for GID^SR4^ [34] are also major ubiquitylation targets of Chelator-GID^SR4^, we performed an activity assay with WT and mutant Mdh2, in which putative target lysine clusters (K254/K256/K259; K330; K360/K361) were mutated to arginines (Fig. S5A). **To** start the reaction, 0.2 µM Uba1, 1 µM Ubc8-6xHis, 0.5 µM E3 GID^SR4^, 0 or 2 µM Gid7, 1 µM WT or mutant Mdh2-6xHis and 20 µM of Ub (WT or all K>R) were mixed. After quenching, Mdh2-6xHis (and its ubiquitylated versions) were visualized by immunoblotting with anti-6xHis primary antibody and HRP-conjugated anti-mouse secondary antibody.

To quantitatively compare recognition of phage display-identified sequences and degrons of natural GID substrates by yGid4, we employed competitive ubiquitylation assays (Fig. 4F). Unlabeled peptide inhibitors comprising the analyzed sequences were titrated to compete off binding of Mdh2-FAM to GID^SR4^, thus attenuating its ubiquitylation. Reactions were started by addition of 20 μM ubiquitin to the mixture of 0.2 μM E1 Uba1, 1 μM E2 Ubc8-6xHis, 0.5 μM E3 GID^Ant^, 1 μM yGid4 (Δ1-115), 0.25 μM of Mdh2-FAM and various concentrations of peptide competitors. After 3 minutes, the reactions were quenched so that they were still in the linear range. The fractions of ubiquitylated Mdh2 in the presence of an inhibitor were divided by that for Mdh2 alone and plotted against peptide concentration. Fitting of the data to [inhibitor] vs. response model in GraphPad Prism yielded IC50 values.

To qualitatively compare degrons of Fbp1 and Mdh2 in the context of full-length substrates (Fig. 6A), we performed activity assay with WT and degron-swapped versions (Fbp1^Mdh2 degron^ and Mdh2^Fbp1 degron^) of the substrates by mixing 0.2 μM E1 Uba1, 1 μM E2 Ubc8-6xHis, 1 μM E3 GID^Ant^, 2 μM yGid4 (Δ1-115), 0.5 μM of WT or mutant version of Fbp1-FAM or Mdh2-FAM and 20 μM ubiquitin.

Kinetic parameters for ubiquitylation of WT and degron-swapped versions of Fbp1 and Mdh2 were determined as described previously [38]. Briefly, to determine Michaelis-Menten constant (*K*_m_), we titrated E3 (GID^SR4^ or Chelator-GID^SR4^) at constant substrate concentration kept below *K*_m_ (0.5 and 0.1 μM for reactions with GID^SR4^ and Chelator-GID^SR4^, respectively; Fig. 5A, 5B, S4E). The reaction time was optimized so that the initial velocity for all reactions was in the linear range. The fraction of ubiquitylated substrate was calculated and plotted as a function of E3 concentration in GraphPad Prism and fit to Michaelis-Menten equation to determine *K*_m_. To calculate k_cat_, time course assays were performed with the ratios of [E3]:*K*_m_ and [substrate]:*K*_m_ kept the same for all substrates and E3 versions (2.7 and 0.4, respectively; Fig. 5C). The rates of the reactions were calculated by linear regression in GraphPad Prism from plots of fraction of ubiquitylated substrates vs. reaction time (Fig. S4D) and converted into initial velocity using the following equation: V_0_ = *rate* · [*substrate*]. Then, V_max_ was estimated using a modified form of the Michaelis-Menten equation: 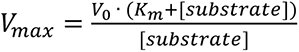. To obtain *k_cat_* values, *V_max_* was divided by the E3 concentration: 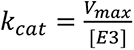.

### Yeast strain construction and growth conditions

The yeast strains used in this study are specified in the Reagents table. All the yeast strains were constructed as derivatives of BY4741 using standard genetic techniques (Knop et al., 1999; Janke et al., 2004; Storici and Resnick, 2006) and were verified using PCR, DNA sequencing and immunoblotting to confirm protein expression.

### *In vivo* yeast substrate degradation assays

In order to test the effect of degron identity on glucose-induced degradation of GID substrates, we monitored turnover of WT and degron-exchanged versions of Mdh2 and Fbp1, using the promoter reference technique adapted from [45]. Initially, WT and ΔGid7 yeast strains were transformed with a plasmid harboring the open reading frame of either Fbp1-3xFLAG, Mdh2-3xFLAG or their mutant versions (Fbp1^Mdh2 degron^-3xFLAG and Mdh2^Fbp1 degron^-3xFLAG) and the control protein DHFR-3xHA, both expressed from identical promoters. Cells were then grown in SD-glucose medium to OD_600_ of 1.0 followed by carbon starvation in SE medium (0.17% yeast nitrogen base, 0.5% ammonium sulfate, 2% ethanol, amino acid mix) for 19 hours. Next, yeast at the equivalent of 1 OD_600_ was transferred to SD-glucose medium containing 0.5 mM tetracycline resulting in translation inhibition induced by its binding to specific RNA-aptamers within ORFs of the examined and control proteins. At the indicated time points, 1 mL of cells were harvested and pellets were flash frozen in liquid nitrogen. Cell lysis was performed by thawing and resuspending the pellets in 800 μL 0.2 M NaOH, followed by 20 min incubation on ice and subsequent centrifugation at 11,200xg for 1 minute at 4°C. The supernatant was removed and pellets were resuspended in 50 μL HU buffer (8 M Urea, 5% SDS, 1 mM EDTA, 100 mM DTT, 200 mM Tris pH 6.8, protease inhibitor, bromophenol blue), heated at 70°C for 10 minutes and then centrifuged again for 5 minutes at 11,200xg and at 4°C. The substrates and the control protein DHFR were visualized by immunoblotting with, respectively, anti-FLAG or anti-HA primary and DyLight fluorophore conjugated secondary antibodies, and imaged using a Typhoon scanner (GE Healthcare). Quantification was done using the ImageStudioLite software (LI-COR). For the final graphs, the substrate signal was first normalized relative to the DHFR signal and then to the time point zero (before glucose replenishment). Three biological replicates were performed for all the assays.

### X-ray crystallography

All crystallization trials were carried out in the MPIB Crystallization facility. All crystals were obtained by vapor diffusion experiment in sitting drops at room temperature.

Crystals of hGid4 (Δ1-99) (without a peptide) were obtained at a concentration of 10 mg/ml using 18% PEG 3350 with 0.2 M ammonium nitrate and 0.1 M Bis-Tris buffer at pH 7. Crystals were cryoprotected in 20% ethylene glycol and flash-frozen in liquid nitrogen for data collection. The diffraction dataset was recorded at beamline PXII, Swiss light Source (SLS) in Villingen, Switzerland.

For hGid4 (Δ1-120, Δ294-300) crystals containing FDVSWFM peptide, 9.2 mg/mL of hGid4 was mixed with 600 μM FDVSWFM peptide and incubated for 1 h on ice before setting up trays. Crystals were obtained using 1.1 M Sodium malonate, 0.3% Jeffamine ED-2001 pH 7 and 0.1 M HEPES pH 7 and cryoprotected using mix of 20% glycerol and 20% ethylene glycol. The diffraction dataset was recorded at beamline PXII, Swiss light Source (SLS) in Villingen, Switzerland.

Similarly, for yGid10 (Δ1-64, Δ285-292) crystals with the peptide FWLPANLW, the protein was concentrated to 10 mg/mL and mixed with the peptide to obtain final protein and peptide concentrations of 262 μM and 760 μM, respectively (∼3-fold molar excess of the peptide). Crystals were obtained using 0.1 M MES pH 6.9 and cryoprotected using 20% ethylene glycol. The diffraction dataset was recorded at X10SA beam line, Swiss Light Source (SLS) in Villingen, Switzerland.

All the crystal data were indexed, integrated, and scaled using XDS package. Phasing was performed through molecular replacement using the previous structure of hGid4 (PDB: 6CCR, in case of hGid4 with and without a peptide) or cryo EM structure of yGid4 (extracted from PDB: 7NS3, in case of peptide-bound yGid10) using PHASER module integrated into PHENIX software suite (Adams et al., 2010; Afonine et al., 2018; DiMaio et al., 2013). Model building was done using Coot (Emsley and Cowtan, 2004; Emsley et al., 2010), and further refinements were carried out with phenix.refine. Details of X-ray diffraction data collection and refinement statistics are listed in Table S1.

## Supplemental information

**Figure S1:**
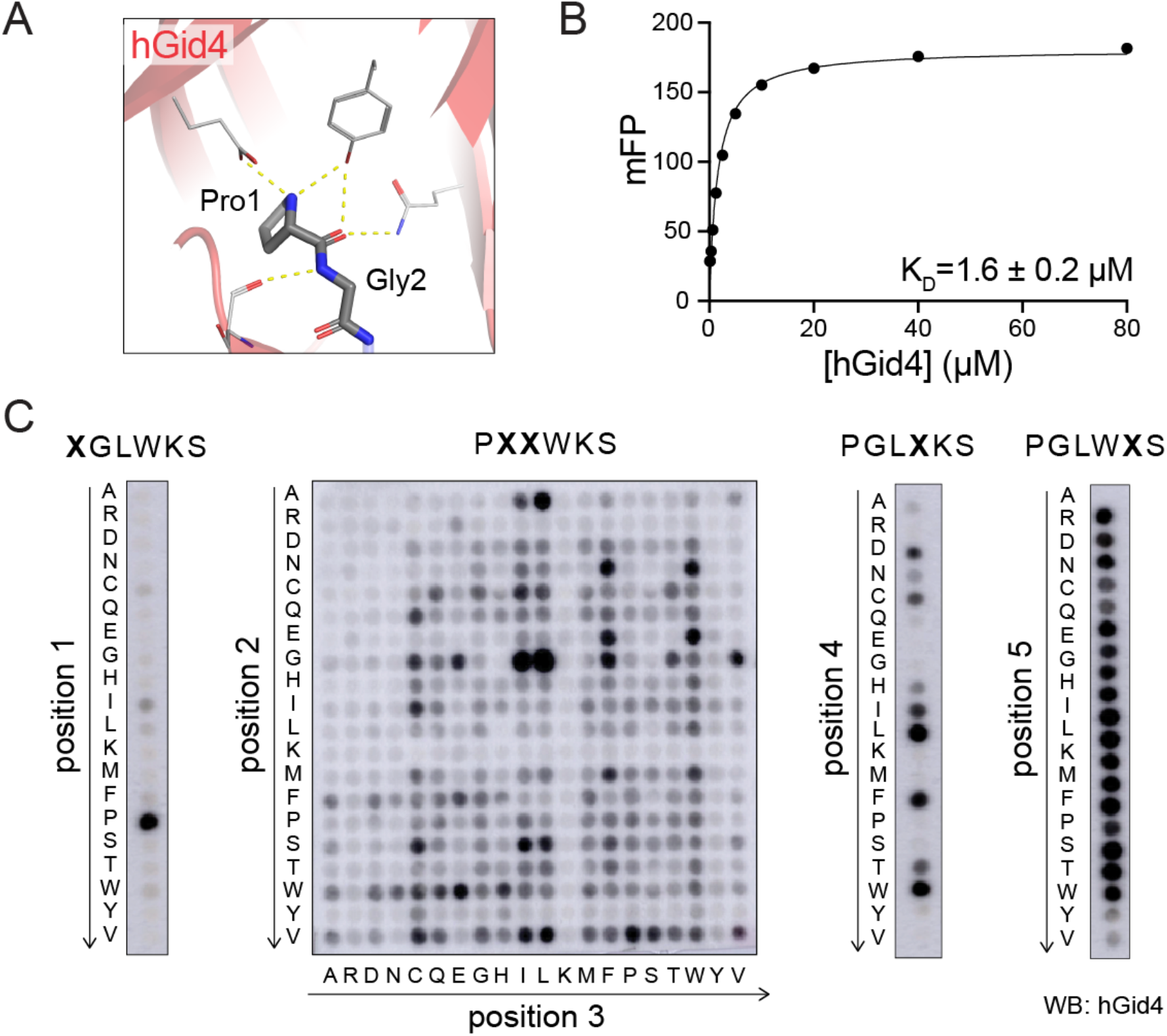
Characterization of hGid4 specificity toward a Pro/N-terminal peptide. A. Close-up of hGid4 interactions with first two residues of the PGLW peptide (PDB: 6CDC) B. Fluorescence polarization (FP) experiment to quantify binding of fluorescent PGLWKS (C-terminally labelled with fluorescein) to hGid4. Fitting the FP values at increasing hGid4 concentrations to one site binding model yielded K_D_. C. Peptide spot arrays to systematically screen the influence of amino acid substitutions at various positions of the PGLWKS sequence on hGid4 binding. Binding of hGid4 to generated peptide derivatives was visualized by immunoblotting with anti-hGid4 antibodies and chemiluminescence. The intensity of the signal corresponds to the binding strength of a given peptide to hGid4.

**Figure S2:**
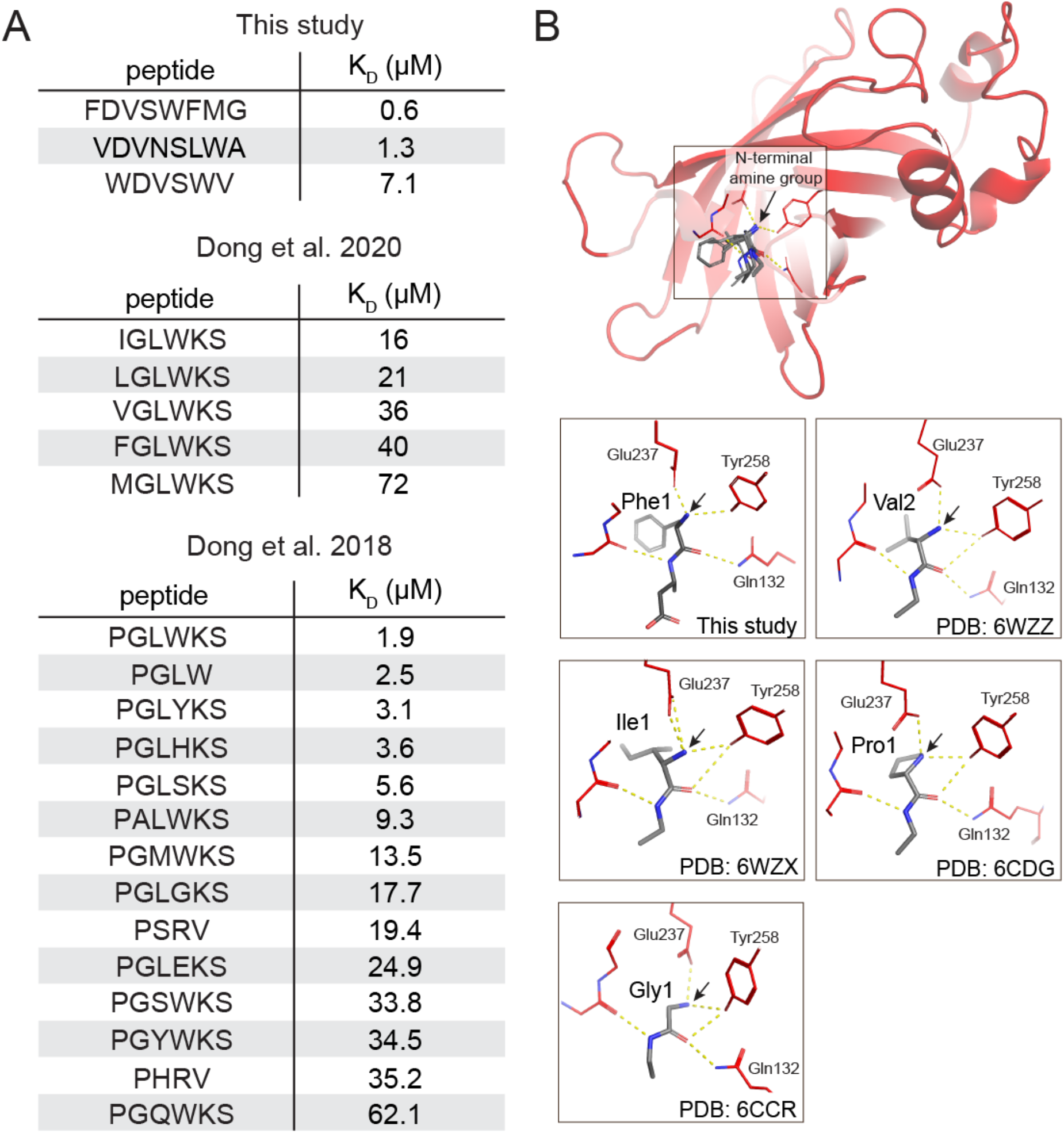
Recognition of diverse peptide sequences by hGid4. A. Table summarizing dissociation constants (K_D_) of various peptides binding to hGid4 measured with ITC in this and the previous studies. B. Overlay of FDVSWFMG-bound hGid4 with the published coordinates of hGid4 (top, first two residues of interacting peptides in all structures shown as grey sticks). The common binding mode of first two N-terminal peptide residues is shown for each structure separately (bottom). Black arrows indicate positions of N-terminal amine groups.

**Figure S3:**
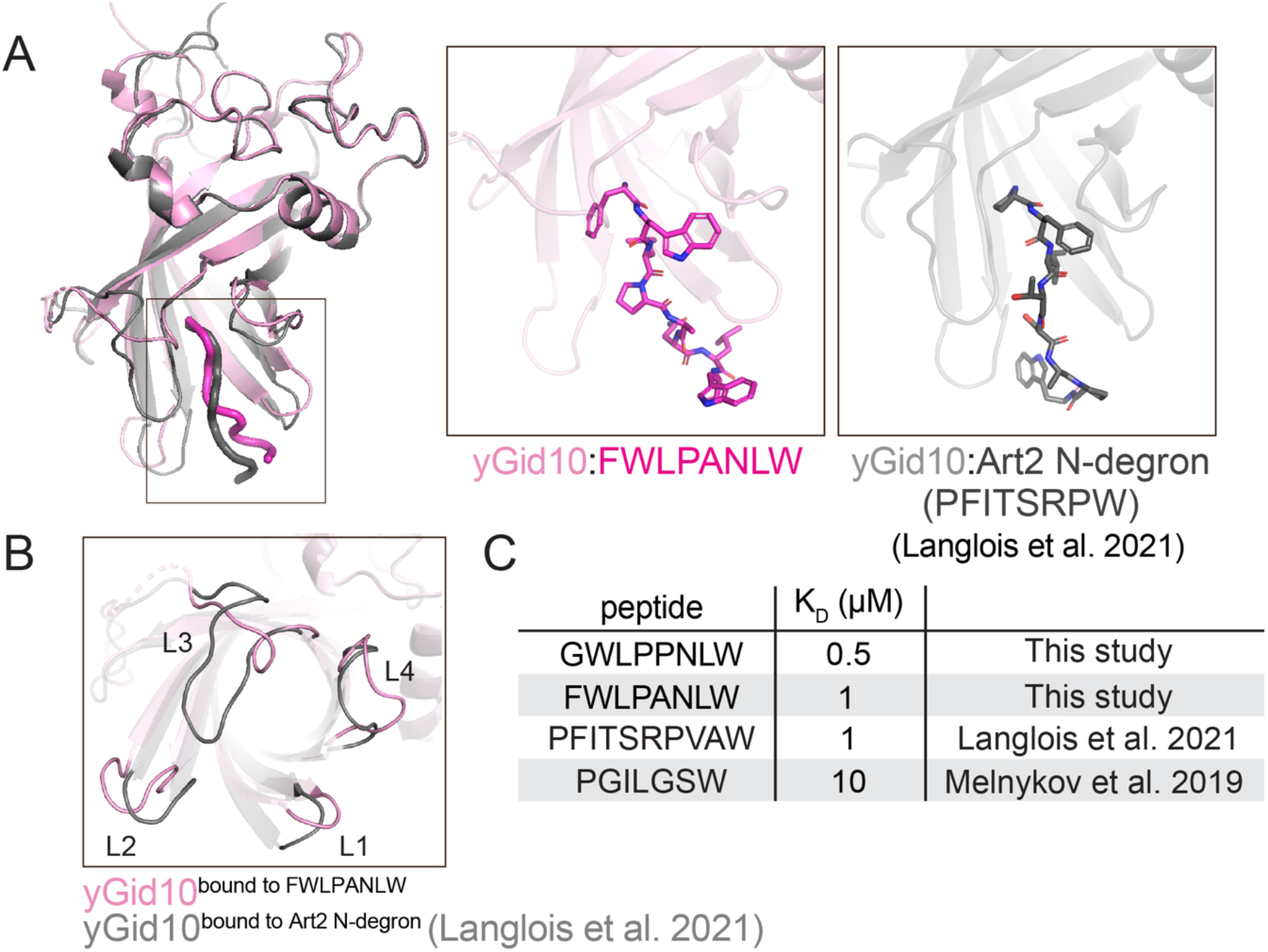
Binding specificity of yGid10. A. Overlay of yGid10 (Δ1-64, Δ285-292) structures (left) bound to FWLPANLW peptide and Pro/N-degron of yGid10 substrate Art2 (Langlois et al. 2021) (shown as magenta and grey ribbons, respectively). Close-ups of both peptides binding to the same yGid10 substrate binding tunnel (middle and right). B. Conformational flexibility of yGid10 substrate-binding loops revealed by superimposing of yGid10 structures bound to FWLPANLW (pink) and Art2 degron (Langlois et al. 2021, grey). C. Table summarizing dissociation constants (K_D_) of various peptides binding to yGid10 measured with ITC.

**Figure S4:**
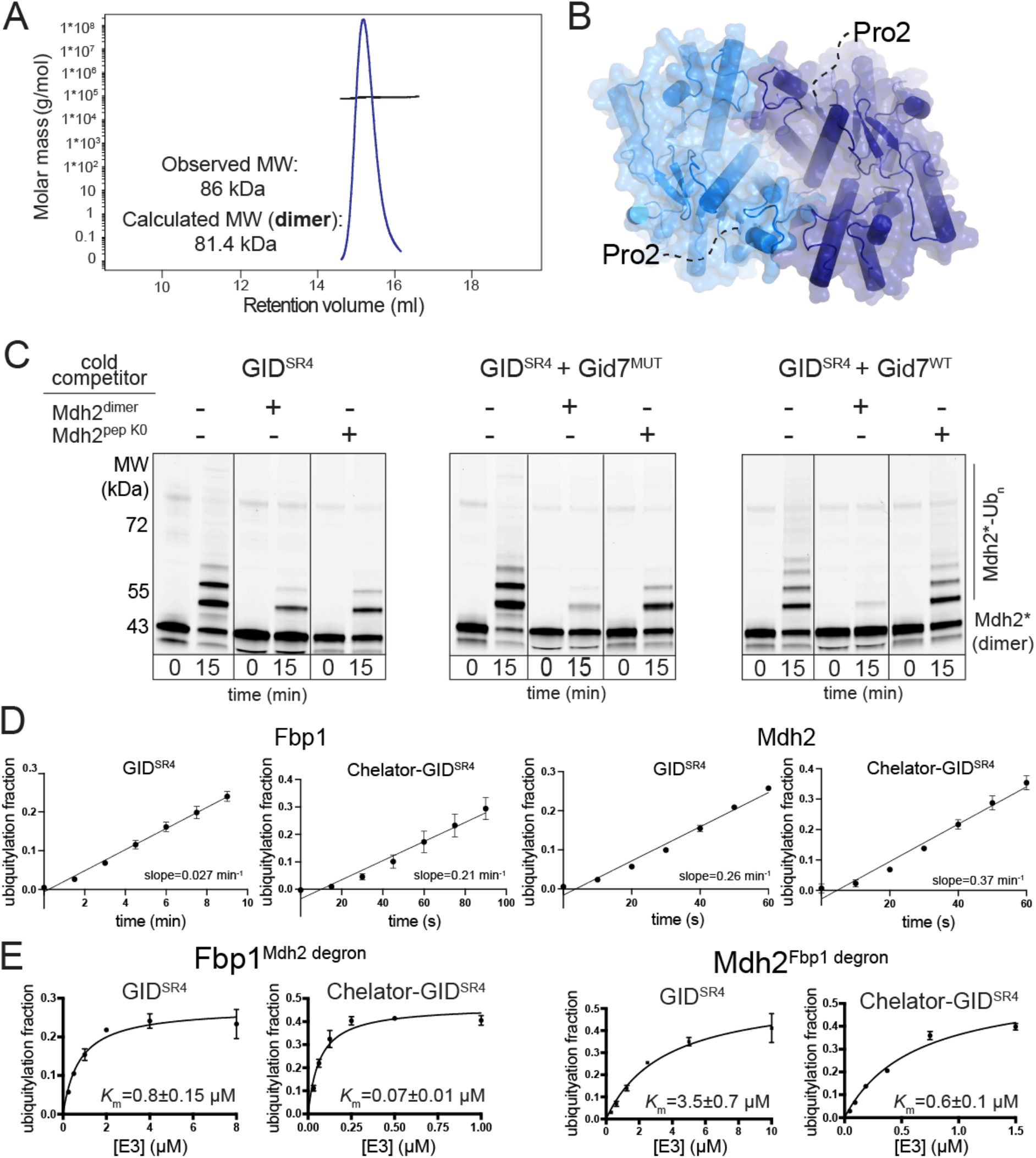
Characterization of native substrate ubiquitylation by GID E3 ligase. A. SEC-MALS analysis of Mdh2 oligomeric state B. Homology model of Mdh2 with unstructured Pro/N-degrons represented as dotted lines C. Competitive ubiquitylation assays probing avid binding of Mdh2 to Chelator-GID^SR4^. Ubiquitylation of fluorescent Mdh2 by monovalent (GID^SR4^ alone or mixed with Gid7^286-745^ mutant) or divalent (Chelator-GID^SR4^) version of GID E3 was competed with monodentate (Mdh2^pep K0^ - lysineless peptide harboring Mdh2 N-terminus) or bidentate (Mdh2^dimer^) inhibitor. D. Time-courses of Fbp1 and Mdh2 ubiquitylation by GID^SR4^ and Chelator-GID^SR4^ employed to determine *k*_cat_ values (Fig. 5C) E. Plots showing fraction of *in vitro*-ubiquitylated degron-swapped Fbp1 and Mdh2 as a function of varying concentrations of GID^SR4^ or Chelator-GID^SR4^. Fitting to Michaelis-Menten equation yielded *K*_m_ values. Error bars represent standard deviation (n=2).

**Figure S5:**
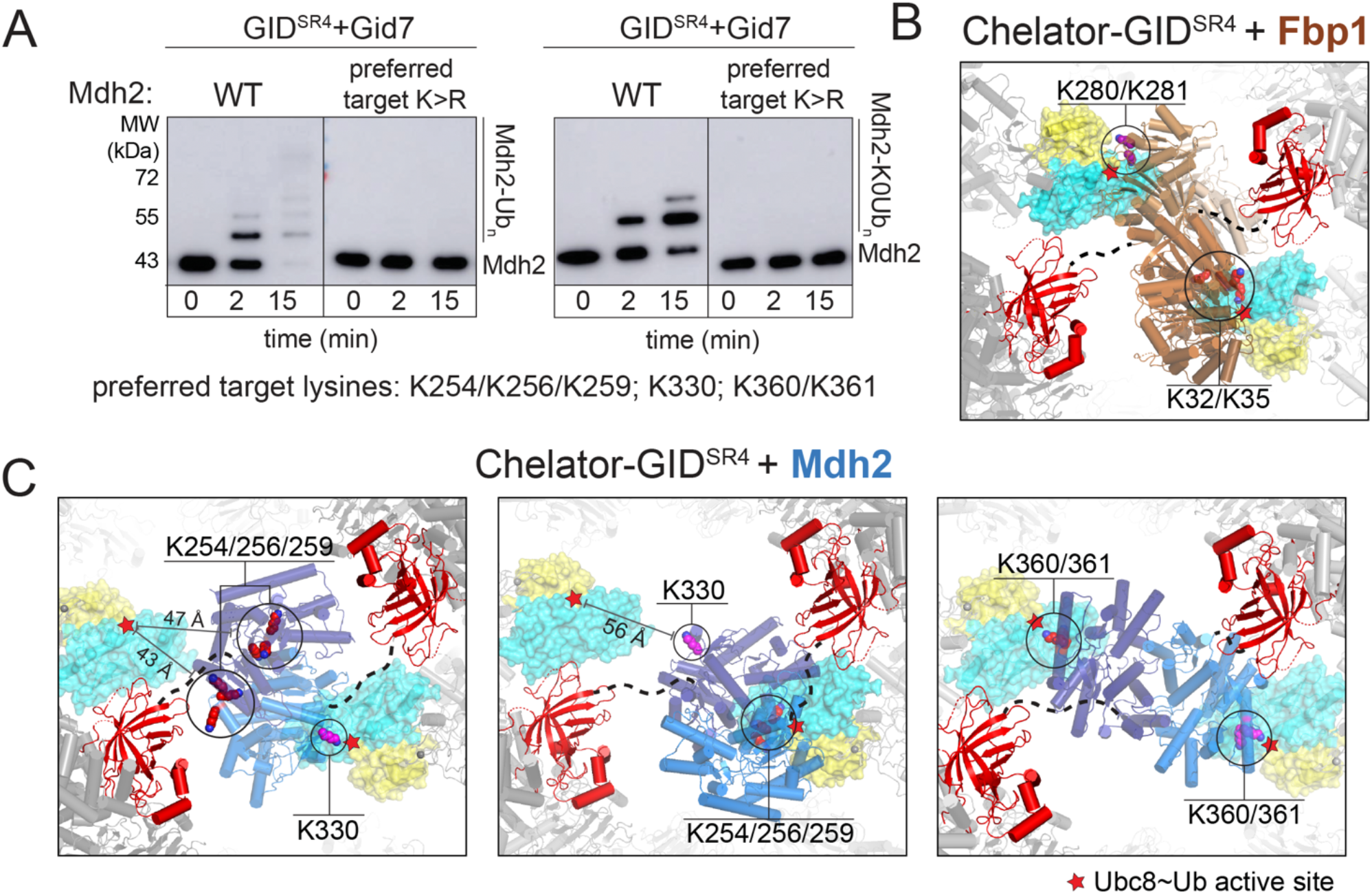
Structural modeling of Fbp1 and Mdh2 ubiquitylation by Chelator-GID^SR4^. A. *In vitro* assay of Mdh2-6xHis testing effect of mutating several of its lysines (previously determined to be preferred targets of GID^SR4^ by mass-spectrometry) on Chelator-GID^SR4^-dependent ubiquitylation. Mdh2-6xHis and its ubiquitylated versions were visualized by anti-6xHis immunoblotting. Reactions were performed with WT and lysine-less ubiquitin (Ub; K0 has all Lys mutated Arg) B. Ubiquitylation model of Fbp1 (PDB: 7NS3, 7NS4, 7NS5, 7NSB; EMD-12557) involving juxtaposition of its target lysines (red and violet sticks, indicated by black circles) with Ubc8∼Ub active site (red stars) generated by: (1) docking of two Fbp1 degrons (black dashes) into substrate binding cavities of two opposing Gid4 molecules (red cartoon) and (2) rotation of the folded Fbp1 domain (brown cartoon) so that its target lysines could simultaneously reach both active sites (the recruited and activated Ubc8∼Ub intermediate shown as cyan (Ubc8) and yellow (Ub) surface) modeled by aligning a previous RING-E2∼Ub structure (PDB: 5H7S) with Gid2 RING (grey cartoon). C. Ubiquitylation model of Mdh2 (blue cartoon; obtained by homology modeling) generated as in (B) but requiring a more pronounced shift of substrate receptor-scaffolding modules (Gid1-Gid8-Gid5-Gid4; grey cartoon) towards the center of the oval chelator assembly to enable capture of two Mdh2 degrons (black dashes). Besides K360/361, Mdh2 cannot be oriented so that its target lysines simultaneously engage both Ubc8∼Ub active sites (red stars).

**Table S1.** X-ray crystallography data collection and refinement statistics. Values for the highest-resolution shell are given in parentheses.

|  | yGid10 +<br>FWLPANLW | hGid4 +<br>FDVSWFMG | hGid4 (without<br>a peptide) |
| --- | --- | --- | --- |
| **Data Collection** |  |  |  |
| Spacegroup | P3<sub>1</sub> 2 1 | P4<sub>1</sub> 3 2 | P6<sub>5</sub> |
| Cell dimensions |  |  |  |
| a, b, c (Å) | 99.23 99.23<br>81.02 | 131.31 131.31<br>131.31 | 72.02 72.02<br>146.32 |
| $\alpha, \beta, \gamma$ (°) | 90 90 120 | 90 90 90 | 90 90 120 |
| Resolution range (Å) | 42.97 - 2.22<br>(2.30 - 2.22) | 43.77 - 3.16<br>(3.27 - 3.16) | 57.38 - 3.08<br>(3.19 - 3.08) |
| *I* / $\sigma$ (*I*) | 16.2 (1.1) | 9.1 (1.0) | 8.8 (1.0) |
| Completeness (%) | 97.3 (76.6) | 99.9 (99.6) | 99.6 (96.2) |
| **Refinement** |  |  |  |
| Refinement program | Phenix<br>version 1.19 | Phenix<br>version 1.19 | Phenix<br>version 1.19 |
| Resolution (Å) | 2.22 | 3.16 | 3.08 |
| R<sub>work</sub> / R<sub>free</sub> | 0.19/0.22 | 0.24/0.29 | 0.24/0.27 |
| Reflections used in<br>refinement | 22531 (1756) | 7072 (690) | 7950 (753) |
| Reflections used for R-free | 1127 (88) | 383 (39) | 397 (38) |
| No. of molecules in ASU | 1 | 1 | 2 |
| Total no. of atoms | 1848 | 1403 | 2742 |
| Protein | 1685 | 1332 | 2710 |
| Peptide | 75 | 62 | 0 |
| Ligand | 1 | 0 | 0 |
| Water | 87 | 9 | 32 |
| Wilson B-factor (Å<sup>2</sup>) | 59.5 | 102.1 | 91.8 |
| RMSD bond length (Å) | 0.004 | 0.005 | 0.002 |
| RMSD bond angle (°) | 0.795 | 0.780 | 0.490 |
| Ramachandran<br>favored (%) | 94.2 | 92.5 | 92.8 |
| Ramachandran<br>allowed (%) | 5.8 | 7.5 | 7.2 |
| Ramachandran<br>outliers (%) | 0 | 0 | 0 |

**Table S2.**
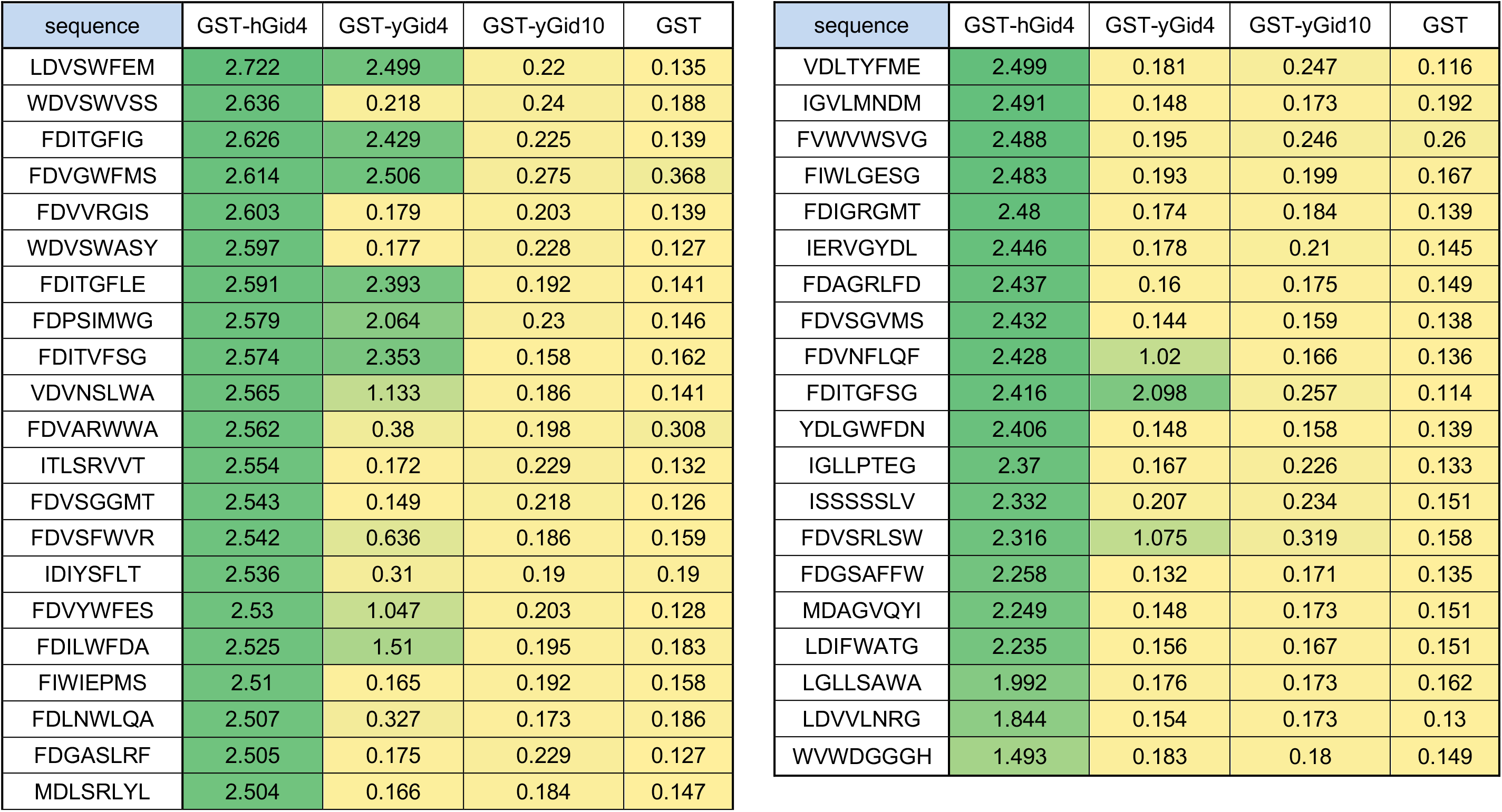
List of phage display-identified sequences binding hGid4 (Δ1-99) and yGid4 (Δ1-115). Sequences were sorted and colored based on increasing intensity of phage ELISA signal for GST-hGid4 (Δ1-99) (from yellow to green; tested also for binding to GST-yGid10 (Δ1-56) and GST-only controls).

**Table S3.** List of phage display-identified sequences binding yGid10 (Δ1-56). Sequences were sorted and colored based on the intensity of phage ELISA signal for GST-yGid10 (Δ1-56) (from yellow to green; tested also for binding to GST-hGid4 (Δ1-99), GST-yGid4 (Δ1-115) and GST-only controls).

| sequence | GST-hGid4 | GST-yGid4 | GST-yGid10 | GST |
| --- | --- | --- | --- | --- |
| FWLPANLS | 0.313 | 0.197 | 2.074 | 0.261 |
| AWLPPNIM | 0.16 | 0.157 | 2.023 | 0.169 |
| GFLPPNLV | 0.208 | 0.189 | 1.911 | 0.146 |
| GWLPQNLM | 0.203 | 0.183 | 1.819 | 0.158 |
| GFLPPNLL | 0.13 | 0.128 | 1.729 | 0.119 |
| GFLPPNLG | 0.173 | 0.178 | 1.723 | 0.174 |
| AWLPPNLE | 0.194 | 0.167 | 1.691 | 0.141 |
| AWLPPNVV | 0.178 | 0.167 | 1.623 | 0.179 |
| GWLPPNFR | 0.203 | 0.178 | 1.263 | 0.147 |
| FWLPPNQL | 0.197 | 0.175 | 1.259 | 0.155 |
| VWLPPNFY | 0.214 | 0.183 | 0.655 | 0.165 |
| ALLPENLR | 0.151 | 0.15 | 0.644 | 0.151 |

